# Effects of serotonin agonists LSD and 25CN-NBOH on conditioned place preference and on synaptic plasticity of VTA dopamine neurons in mice

**DOI:** 10.1101/2024.12.12.628157

**Authors:** Lauri V. Elsilä, Elina Nagaeva, Jari-Pekka Luukkonen, Esa R. Korpi

## Abstract

The current research on psychedelic compounds such as lysergic amide diethylamide (LSD) is leaning heavily on the notion that psychedelics are not addictive. While much of the literature supports this argument, some of the common use patterns and the descriptions of the subjective effects of these compounds in humans, together with rather lacking and mixed data from non-human animal studies leave room for questions of potentially rewarding or reinforcing stimulus effects. Initiated by a surprising finding in a control study, we investigated these potential rewarding effects of LSD and a selective 5-HT_2A_ agonist 25CN-NBOH using both unbiased and biased designs of conditioned place preference as well as *ex vivo* patch-clamp electrophysiology measurements of glutamatergic synaptic plasticity on midbrain ventral tegmental area (VTA) dopamine neurons in C57Bl6/J mice. Our results showed no reliable formation of place preference with either compound, agreeing with previous claims of psychedelics having at most weak reinforcing effects. However, we did observe single doses of the drugs, especially LSD, inducing synaptic plasticity in the medially located VTA dopamine neurons, implicating a role for the midbrain dopamine system in the effects of psychedelic drugs.

**Graphical abstract:** Treatment with mixed serotonin receptor agonist, psychedelics lysergic acid diethylamide (LSD) or selective serotonin 2A receptor agonist 25CN-NBOH did not cause reliable induction of conditioned place preference in C57Bl/6J mice. However, we did observe single doses of the drugs, especially LSD, inducing synaptic plasticity in the medially located VTA dopamine neurons. These findings challenge some of the previous rodent data but are in general in line with the claims of psychedelics having at most weak reinforcing effects.

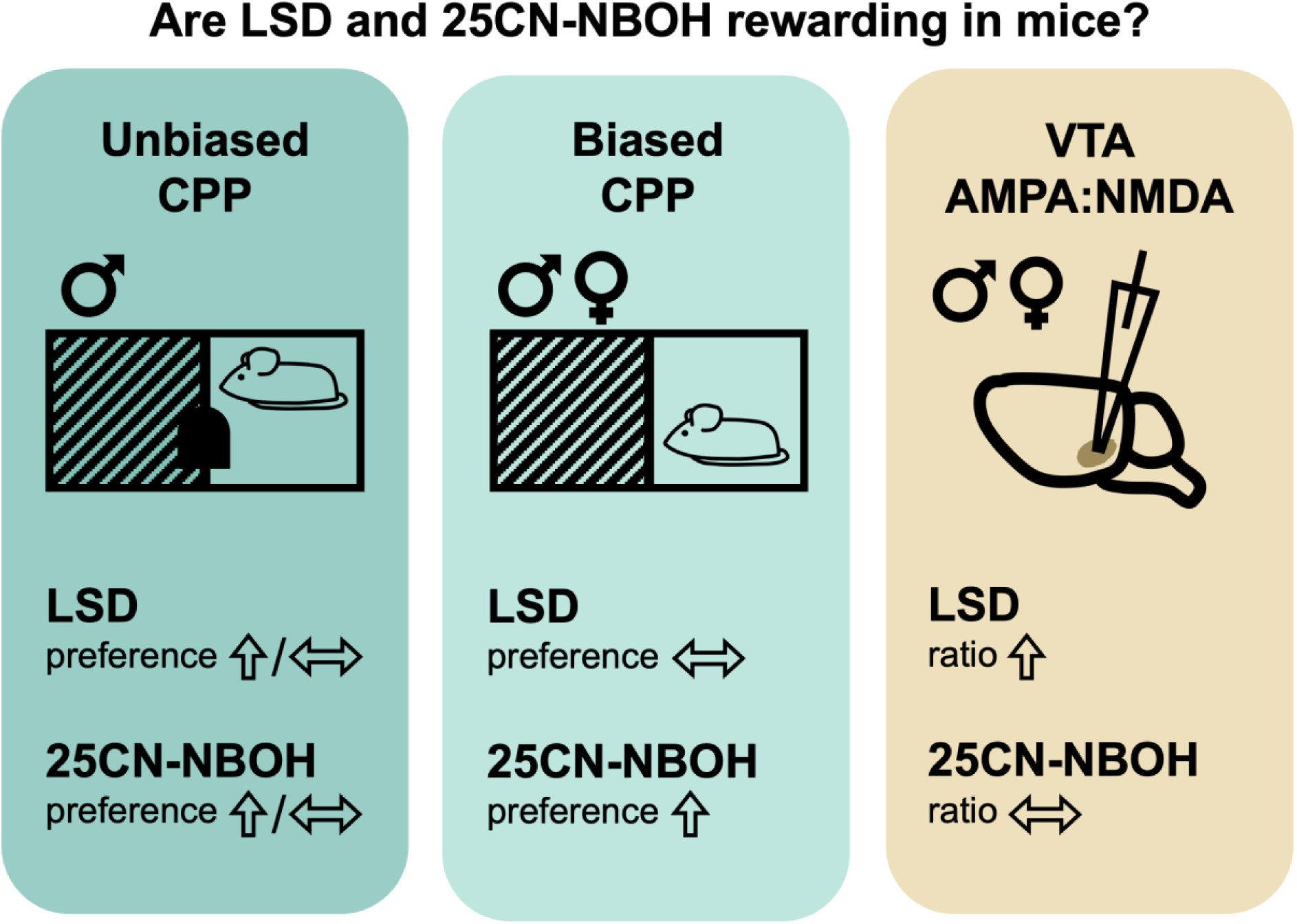

## 1 INTRODUCTION

Renewed interest in psychedelic drugs, especially in the mixed serotonin receptor agonists (i.e. classical psychedelics) such as lysergic acid diethylamide (LSD), psilocybin, and N,N-dimethyltryptamine (DMT), has also fortified the common notion that psychedelics are not addictive (e.g. Nichols, 2016; Grieco *et al*., 2022; Nutt *et al*., 2023). Despite the classifications for hallucinogen use disorders in the major diagnostic systems ICD-11 and DSM5, there are only singular clinical reports of patients describing themselves as being dependent on psychedelics (Modak *et al*., 2019) whereas the prevailing opinion in the literature is that there is either no or very weak evidence of abuse potential for psychedelics (Amsterdam *et al*., 2011; Johnson *et al*., 2018; Schlag *et al*., 2022; Konradi & Hurd, 2023). On the other hand, different studies show that there are people that tend to use psychedelics repeatedly and who report to use psychedelics for fun (Basedow & Kuitunen-Paul, 2022; Glynos *et al*., 2023). People also often describe the subjective effects of psychedelics to include generally positive qualities (drug liking), increased mood, feelings of bliss, ecstasy, and euphoria (although notably also feelings of dread and anxiety; Schmid *et al*., 2015; Preller & Vollenweider, 2018; Carbonaro *et al*., 2020; Holze *et al*., 2022; Ley *et al*., 2023). Together, these might point towards psychedelics having some hedonic, rewarding, or reinforcing stimulus properties.

The claim of non-addictiveness is commonly paired with evidence that non-human animals do not self-administer psychedelics, which seems to be mostly true for non-human primates (Hoffmeister, 1975; Griffiths *et al*., 1979; Yanagita, 1986, but see also Siegel & Jarvik, 1980; Fantegrossi *et al*., 2004; Goodwin, 2016). To our knowledge, there are no published data on whether this holds true for rodents. The self-administration literature is well supported by findings from rodent intracranial self-stimulation studies where LSD, psilocybin, mescaline, a 5-HT_2A/2C_ agonist 2,5-dimethoxy-4-iodoamphetamine (DOI), and a selective 5-HT_2A_ agonist (4-bromo-3,6-dimethoxybenzocyclobuten-1-yl)methylamine (TCB-2) did not induce reinforcing effects, but, on the contrary, potentially slightly aversive effects (Katsidoni *et al*., 2011; Sakloth *et al*., 2019; Elsilä *et al*., 2022; Jaster *et al*., 2022). However, these findings are contrasted by studies showing that LSD (Parker, 1996; Meehan & Schechter, 1998) and semi-synthetic pro-drug of psilocin, 4-acetoxy-N,N-dimethyltryptamine (4-AcO-DMT; Vargas-Perez *et al*., 2024) can cause conditioned place preference in rats; similar findings have not been published for mice. The evidence base from the preclinical studies is therefore mixed and lacking, especially for mice, the most used animal species in the preclinical models.

The literature on drug discrimination proves that psychedelics have interoceptive stimulus properties in non-human animals (reviewed in Baker, 2018), but as noted for example by Johnson and Ettinger (2000), discriminative stimulus effects do not automatically mean reinforcing stimulus effects. Methods like conditioned place preference (CPP) can then be used to further elucidate the stimulus properties. CPP is a paradigm based on Pavlovian conditioning where an originally neutral stimulus (conditioned stimulus, CS) evokes an approach behaviour (conditioned response, CR) after being paired with a motivationally significant experience (unconditioned stimulus, US; Hall, 1994; Tzschentke, 1998). It is often used in preclinical addiction research to investigate the motivational properties of drugs (Cunningham *et al*., 2011; Koob *et al*., 2014) by using the drug of interest as the US (Tzschentke, 1998). The ability of the CS to elicit approach behaviour during the test session after the conditioning is commonly considered to mean that the US, the drug, has positively reinforcing stimulus properties.

The mesolimbic dopamine system plays an important role in conveying motivationally salient stimuli into adaptive behavioural responses (Lammel *et al*., 2014). These stimuli not only cause complex activation patterns in the dopamine system (Berridge *et al*., 2009; Volkow *et al*., 2017; Klawonn & Malenka, 2018) but are also known to induce plastic changes in different parts of the system (Thomas & Malenka, 2003; Lüscher & Malenka, 2011). One of the most commonly used experimental measures for synaptic strength is the ratio between the excitatory post synaptic currents mediated by the α-amino-3-hydroxy-5-methyl-4-isoxazolepropionic acid (AMPA) receptors and the N-methyl-D-aspartic acid (NMDA) receptors, also known as the AMPA:NMDA ratio (Lüscher & Malenka, 2011). It is a commonly used physiological readout in preclinical addiction research as most drugs of abuse are known to produce a robust and long-lasting increase in the ratio in the midbrain ventral tegmental are (VTA) dopamine neurons 24 h after only a single dose of the drug. These plastic changes have also been observed after self-administration of and Pavlovian conditioning with palatable food (Chen *et al*., 2008; Stuber *et al*., 2008), as well as also after exposure to aversive experiences (Lammel *et al*., 2011; de Jong *et al*., 2019), and together with behavioural readouts this can be a useful tool in further explicating the salient quality of any drug stimuli.

Here, we report on a set of experiments in mice investigating whether a classical psychedelic, a mixed serotonin receptor agonist LSD, or a selective serotonin receptor 2A agonist N-(2-hydroxybenzyl)-2,5-dimethoxy-4-cyanophenylethyl-amine (25CN-NBOH) would show evidence of inducing positive reinforcement comparable to other drugs of abuse, such as morphine, by studying CPP behaviour and glutamate receptor synaptic plasticity in the VTA dopamine neurons in mice. Originally inspired by a surprise finding, described here as the experiment CPP1, indicating that 25CN-NBOH potentially promoted CPP, the question was further explored with a comprehensive set of CPP experiments. Parallel to these, dopamine neuron electrophysiology *ex vivo* in VTA slices was conducted to look for neuronal adaptations caused by a single drug administration. Based on these experiments we propose that neither LSD not 25CN-NBOH reliably produced place conditioning in C57B/6J mice, although they both showed some plastic changes in glutamatergic synaptic inputs to dopamine neurons located medially in the VTA.

## 2 MATERIALS AND METHODS

All the used materials have been listed with identifiers in the Key Resources Table on the Supplementary Material Table S1.

### 2.1 Study subjects, housing, and handling

The results of this study are based on the data from a total of 224 mice. The conditioned place preference behaviour was studied with C57BL/6JCrl (JAX® Mice Strain, Charles River, Wilmington, MA, USA) male (n=156) and female (n=32) mice, approximately 8–10 weeks of age at the time of the experimentation, received straight from the supplier, and housed in groups of 4 in reversed 12:12 h light cycle (lights off at 06.00 am). The electrophysiological studies were conducted with transgenic mice expressing the enhanced green fluorescent protein under the tyrosine hydroxylase promoter (TH-EGFP; MMRRC no. 000292-UNC, background C57BL/6; Gong et al. 2003), both males (n=19) and females (n=17), approximately 20 days (between 16–23) of age at the time of the experimentation, bred in-house and housed with their same sex littermates in groups of 1–6 in regular 12:12 h light cycle (lights on at 06.00 am). All mice were housed in individually ventilated home cages (GM500, Tecniplast, Buguggiate, Italy) with aspen bedding, in-cage plastic shelters and tunnels, nesting material, wooden blocks, and water and basic rodent chow constantly available. All handling was done either with cupping or tunnel methods (Hurst & West, 2010; Gouveia & Hurst, 2017). This work was carried out with the support of HiLIFE Laboratory Animal Centre Core Facility at the University of Helsinki, and the experiments were approved by the Finnish Project Authorisation Board (permissions no. ESAVI/1172/04.10.07/2018 and ESAVI/1218/2021) and conducted in accordance with national and EU-level ethical and procedural guidelines and legislation.

### 2.2 Habituation

The mice that went through behavioural procedures were accustomed to experimenters’ handling with a 5-day habituation routine before the start of the experiments, as described before (Elsilä *et al*., 2020, 2022). Shortly, the mice were exposed to intensifying handling, starting with the experimenter slowly moving a hand in the cage and only slightly touching the mice and ending with lifting the mice away from the cage with an open palm and letting them freely explore the length of the experimenter’s arm. The mice were also accustomed to the grip needed for the injections. The procedure was performed systematically and resulted in the same minimum level of habituation for all mice prior to starting the experiment. The mice participating in the biased conditioned place preference (see 2.4.2) were also habituated for being handled and injected on Vetbed fabric blankets. The mice tested with electrophysiology methods (2.4.3) were not habituated for handling.

### 2.3 Pharmacological agents and interventions

All the drugs, mixed serotonin receptor agonist lysergic acid diethylamide (LSD; Sigma-Aldrich, St. Louis, MO, USA for CPP1–CPP4 and electrophysiology; and LGC Standards, Wesel, Germany, for CPP5–6), selective serotonin 2A receptor agonist N-(2-hydroxybenzyl)-2,5-dimethoxy-4-cyanophenylethyl-amine hydrochloride (25CN-NBOH, Jensen et al., 2017; a kind gift from prof. Jesper L. Kristensen, University of Copenhagen) and opioid receptor agonist morphine hydrochloride (Yliopiston Apteekki, Helsinki, Finland) were freshly dissolved in sterile physiological saline for injections. The drugs were administered either by intraperitoneal (i.p.) or subcutaneous (s.c.) injections. The used administration route is described separately for every experiment. Injection volume of 10 ml/kg was always used. In each experiment, the investigators administering the drugs, conducting the experiment, and performing the initial data analysis were blind to the treatment groups. The doses for LSD (0.1 and 0.2 mg/kg) and 25CN-NBOH (1.5 and 3.0 mg/kg) were chosen to induce behavioural changes (i.e. ‘hallucinogenic’ doses), using the head twitch response as a proxy measure (Halberstadt *et al*., 2020); responses were not measured specifically for this work, but the doses were based on literature (Halberstadt & Geyer, 2013; Fantegrossi *et al*., 2015; Buchborn *et al*., 2018) and previous work in the laboratory (Elsilä *et al*., 2020). Also, the timings for the behavioural tests were planned so that these subjective effects of the drugs would take place during the conditioning trials. Morphine was used as a positive control with a dose (10 mg/kg) known to produce conditioned place preference (Aitta-aho *et al*., 2012) and increased AMPA:NMDA current ratios in the earlier works of the laboratory (Vashchinkina *et al*., 2018). The doses for 25CN-NBOH and morphine were both calculated in the salt form.

### 2.4 Conditioned place preference

The ability of the drugs to induce conditioned place preference was investigated using two different experimental designs, unbiased and biased ones. All behavioural experiments were conducted during the active phase of the animals, either between 09.00 am and 01.00 pm (CPP1–4), or during the periods of 09.00 am–12.00 pm and 03.00 pm–06.00 pm (CPP5–6).

#### 2.4.1 Unbiased design

Four conditioned place preference experiments (CPP1–4) were done using a forced-choice, unbiased, counterbalanced design based on the description by Cunningham et al. (2006) and a prior study using the same apparatus (Kopra *et al*., 2018). Only male mice were tested with this design (total n=124). The experiments were conducted using infrared beam-based open field activity chambers (ENV-51S; Med Associates, Inc., St. Albans, GA, USA) with a two-chamber place preference inserts (ENV-517; Med Associates). The chambers had different floor materials with different tactile cues and were separable with a guillotine door. One chamber had the original metal grid floor (11×11 mm; Grid) and the other had a custom-made, grey perforated plastic floor insert with round 7.5 mm holes 16 mm apart (Hole). The movement and the location of the animal were recorded by interruption of the infrared beams and analysed with the Activity Monitor software (v. 6.02; Med Associates). The apparatus bias assessment is shown in the Supplementary Material Table S1 and Figure S1. Before and between the experiments, the chambers were thoroughly washed with water and soap, but during the experiments only the visible urine and faecal boli were removed after every session.

The experiments consisted of three phases: habituation, conditioning phase, and preference/post-test phases. In the habituation phase, the mice were placed into the test arena and were let to freely explore both chambers with the door open for 15 min. No injections were given in this phase. In the conditioning phase, the mice received a total of four injections of the test drug (LSD, 25CN-NBOH, saline, or morphine, i.p.; CS+ trials) and four injections of saline (i.p.; CS– trials) on alternating days with 24 h between the consecutive trials. The treatment groups were counterbalanced for the conditioned stimulus so that half of the mice received the drug treatment on the metal grid floor and the other half on the plastic floor, and for the starting order with half of the mice starting with the drug and half with the saline treatment. Immediately after the injection, the mice were placed into the chambers with the guillotine door closed for 30 min. The formation of preference was assessed 24 h after the last conditioning trial with a 15-min preference test, where the mice were placed into the chamber with the door open like in the habituation phase.

The primary outcome measure for the preference formation was the difference between the floor subgroups (Grid and Hole) in the time spent as seconds in the Grid chamber divided by the total duration of the session in minutes (s/min, Cunningham et al., 2003; Table 1); the transformation variable gains values between 0 and 60 with 0 equalling to complete aversion to the target chamber, 60 to complete preference, and 30 as the neutral midpoint, and enables comparison between sessions of different lengths. As secondary outcome measures, the time spent on the CS+ chamber during the habituation and the test session, and the paired difference between them (i.e. time-shift) were also assessed. As the closing of the guillotine door caused an erroneous signal in some of the used chambers, the locomotor activity data for the conditioning days was not considered reliable and was not used for analysis (Figure S4).

**Table 1.** The primary and secondary outcome measures and dependent variables in the conditioned place preference experiments. The evidence of preference is based on Cunningham et al. (2003). All the time scores and time-shifts were measured as s/min transformation units as explained in 2.4.1.

| Design | Experiments | Primary outcome variable | Primary evidence of preference | Secondary outcome variable | Secondary evidence of preference |
| --- | --- | --- | --- | --- | --- |
| ***Unbiased*** | CPP1–4 | Time scores on Grid floor | Difference between the subgroups within each drug treatment | Time-shift within the treatment group | Difference between the Test CS+ and Habituation CS+ within the group |
| ***Biased*** | CPP5–6 | Time-shift, i.e. Test CS+ *minus* Pre-test CS+ | Difference between the drug and the saline group | Time-shift within the treatment group | Difference between the Test CS+ and the Pre-test CS+ |

In the CPP1, the doses 0.2 mg/kg of LSD and 3.0 mg/kg of 25CN-NBOH were tested against saline as a negative control (n=10/group), and CPP2 was a replication of this (n=10/group) conducted by another investigator. To further assess dose-response relationships and to include a positive control, the CPP3 and CPP4 tested doses 0.1 and 0.2 mg/kg of LSD and doses 1.5 and 3.0 mg/kg of 25CN-NBOH, respectively, against saline and 10 mg/kg of morphine (n=8/group in both experiments).

#### 2.4.2 Biased design

To assess the generalisability of the place preference effects, two experiments (CPP5–6) were conducted with another apparatus using a biased, forced choice design with both male and female (n=32/sex) mice (apparatus bias assessment Table S1, Figures S2, and S3). Based on our previous designs (Siivonen *et al*., 2018; Vashchinkina *et al*., 2018), the test was performed in eight polycarbonate cages (TP1290, Tecniplast) arranged in two rows of four. The floors were covered with either of the two floor materials acting as the conditioned stimuli, pink polyethene plastic with 1.2-cm bars separated by a 0.5-cm gap, and blue lattice-like nitrile rubber floor with 1-cm diameter holes. The cages were covered with perforated transparent plastic covers during the trials. The movement and the location of each mouse were recorded with an overhead camera (acA1300-60gm, Basler, Ahrensburg, Germany) connected to Ethovison XT video tracking system (v. 10.1; Noldus, Wageningen, the Netherlands).

Like the unbiased design, these experiments comprised of three phases. In the pre-test phase, the mice were habituated to the arenas and their initial preferences for the used materials were assessed by placing them to the arenas covered in 1:1 ratio with both floor materials for 15 min once in the morning and again in the afternoon. The floor positions (i.e. on which side of the cage which material was placed) were counterbalanced within each trial. The time spent on the floor materials on the second pre-test session was used to determine the preferred and non-preferred material for each mouse. In the conditioning phase, the CS– trials, with saline paired with the preferred material, were conducted in the morning (8–11 am), and the CS+ trials, with the drugs (LSD, 25CN-NBOH, morphine, or saline) paired with the non-preferred material, in the afternoons (3–6 pm). The mice were injected s.c. on a Vetbed fabric blankets and placed to the arenas covered fully with the corresponding floor material for 30 min. The two conditioning sessions were repeated four times on consecutive days. After the last conditioning day, the formation of preference was tested in a single test the following morning with a 15-min session identical to the pre-tests.

The main outcome measure determining the formation of preference was the difference in the time spent on the CS+ material between the test session and the second pre-test session (time-shift; s/min) and its difference from the negative control saline (Table 1). As a secondary outcome measure, the time-shift within each treatment group (i.e. difference between the test and the pre-test) was assessed. The distance moved was also determined for all sessions as a measure of locomotor activity.

Males and females were never tested at the same time but in separate, single-sex sessions, with the starting order of each sex alternating every day. Distinct batches of polycarbonate arenas were used for each sex, but the floor materials were shared with all mice; both the arenas and the floor materials were thoroughly washed with water and diluted odourless dish soap before every use.

The same dose-response set-up as in CPP3 and 4 was used here, with LSD (0.1 and 0.2 mg/kg) and 25CN-NBOH (1.5 and 3.0 mg/kg) being tested against saline and 10 mg/kg morphine in CPP 5 and 6, respectively (n=4/group/sex in each experiment). The treatment groups were balanced between each cage, but within a single cage the treatment group assignments were fully randomised (random number generator and ranking functions in Microsoft Excel).

### 2.5 Electrophysiological measurements

Electrophysiological experiments were performed 24 h after a single intraperitoneal injection of the drug (10 mg/kg morphine n=3, 0.1 mg/kg LSD n=11, or 3.0 mg/kg 25CN-NBOH n=7) or saline. Horizontal 225-μm midbrain slices were prepared as described previously (Nagaeva *et al*., 2021). Shortly, the mice were decapitated, and the brains were dissected in ice-cold sucrose-based cutting solution, containing (in mM): 60 NaCl, 2 KCl, 8 MgCl2, 0.3 CaCl2, 1.25 NaH2PO4, 30 NaHCO3, 10 D-glucose and 140 sucrose, using a vibratome (HM650V, Thermo Scientific, Waltham, MA). After a 15-min incubation at 33°C in artificial cerebrospinal fluid (ACSF) buffer solution (composition in mM: 126 NaCl, 1.6 KCl, 1.2 MgCl2, 1.2 NaH2PO4, 18 NaHCO3, 2.5 CaCl2 and 11 D-glucose), the slices were transferred to room temperature ASCF and kept there during the recordings (4 h).

Measurement of synaptic responses was performed after a 1-h incubation using the whole-cell patch-clamp technique. The currents were amplified (Multiclamp 700A, Molecular Devices, Sunnyvale, CA, USA) and digitized at 20 kHz (Molecular Devices). Glass electrodes (3-4 MΩ) contained (in mM) 130 caesium methanesulfonate, 10 HEPES, 0.5 EGTA, 8 NaCl, 5 QX314, 4 MgATP, 0.3 MgGTP, and 10 BAPTA, pH adjusted to 7.2-7.25 and osmolarity to 280 mOsm. The access and membrane resistances were monitored throughout the experiment. Recordings were discarded if the access resistance changed 20% during the experiment. α-Amino-3-hydroxy-5-methyl-4-isoxazole propionic acid (AMPA) and N-methyl-D-aspartate (NMDA) receptor-mediated excitatory postsynaptic currents (EPSCs) were induced by stimulating glutamatergic afferents every 15 s using a unipolar stimulus glass electrode. The VTA dopamine cells were detected based on green fluorescent protein expressed under tyrosine hydroxylase promotor. For the anatomical division, the cells with relatively larger bodies located near the mammillothalamic tract and with the highest concentration in ventral slices (-4.72 mm and -4.56 mm on dorsal-ventral axis relative to bregma; Franklin & Paxinos, 2008) were defined as the ‘lateral population’ whereas the cells with smaller cell bodies and with the highest concentration near the midline, located in more dorsal slices (-4.56 mm, -4.44 mm, and -4.28 mm relative to bregma) corresponded to the ‘medial population’. The neurons were clamped at +40 mV under blockade of GABA_A_ receptors (picrotoxin, 100 μM), and evoked EPSCs were recorded for 10 min before and after the application of the NMDA receptor blocker D-(-)-2-amino-5-phosphonopentanoic acid (AP5, 50 μM). The AMPA:NMDA ratio was determined by dividing the peak amplitude of the AMPA receptor current by that of the NMDA receptor current. The AMPA receptor current was averaged from 30 EPSCs recorded in the presence of AP5. The NMDA receptor current component was then calculated by subtracting the AMPA current from the total current recorded in the absence of AP5.

### 2.6 Data analysis

Both null hypothesis significance testing and estimation statistics approaches were used for the data analysis. All the statistical hypothesis testing was conducted with InVivoStat software (v. 4.9; Bate and Clark, 2014; Clark et al., 2012) and the related graphs were produced with Prism software (v.10.1.2, GraphPad Software, Boston, MA, USA). The estimation statistics and the related graphs were produced with the Estimation Statistics web application (https://www.estimationstats.com; Ho et al., 2019). All the data are presented as means with the 95% confidence intervals or as individual data points (Cumming, 2014; Cumming & Calin-Jageman, 2017). In the experiments using morphine as a positive control, the data from the morphine group was not included in the main statistical model assessing the effects of LSD or 25CN-NBOH, but a separate test against saline control was used to evaluate the results of the experiment (two-way analysis of variance in CPP3–4, unpaired t-tests in CPP5–6 and electrophysiology, and two-way repeated measures mixed model analysis for the locomotor data).

For each parametric test, the assumptions of the homogeneity of variance and the normal distribution of the residuals were assessed with the predicted vs. residual plots and normal probability plots, respectively (Bate & Clark, 2014). The data from the electrophysiological experiments, violating the assumptions, were log_10_ transformed prior to the analysis to stabilise the variance.

The data from the unbiased conditioned place preference tests (CPP1–4) were analysed using a two-way analysis of variance (ANOVA) approach, with Floor and Treatment as the independent factors and the time on Grid (in s/min) as the main dependent outcome variable. The main analysis was followed by all-pairwise comparisons between the predicted means of the Floor × Treatment interaction using Holm’s procedure to control the family-wise error rate. When analysing the combined data from CPP1 and CPP2, Experiment was used as a blocking factor with two levels.

The data from the biased conditioned place preference designs (CPP5–6) were analysed using one-way ANOVA with Treatment as the main independent variable, the time-shift (time on the drug-paired floor during the test *minus* the pre-test; s/min) as the dependent outcome variable, and Sex as a blocking factor with two levels. This was followed up with pairwise comparisons of the Treatment factor against the saline control group mean using Holm’s correction.

The locomotor activity data from the experiments CPP5 and CPP6 were analysed using a two-way repeated measures mixed model approach, with the distance moved during the 30-min conditioning sessions as the dependent outcome variable (in m), Treatment as the independent variable, Session as the repeated factor and Sex as a blocking factor. This was followed by planned comparisons with Holm’s correction on the predicted means to compare the levels of the effect Treatment × Session.

The electrophysiological data were analysed as AMPA:NMDA ratios after the log_10_ transformation, using one-way ANOVA. Treatment was used as the independent variable and Sex as a blocking factor. Follow-up comparisons of the Treatment factor back to the saline control group were also corrected with the Holm’s procedure. The data were analysed both as average ratios (using the average of 1 or 2 cells measured per animal) and as single ratio values per animal when analysing the data based on the anatomical division. The data from morphine treated mice were not included in the lateral-medial division analysis due to the small number of datapoints per anatomical site.

In addition to the hypothesis testing, estimation based on confidence intervals was introduced by plotting the relevant paired (CPP1–4) or unpaired (CPP5–6, electrophysiology) mean differences with Cumming plots, based on bootstrapped sampling distributions with 5000 bootstrap samples using bias corrected and accelerated 95% confidence intervals (Ho *et al*., 2019). Additionally, the changes in the secondary outcome measures of the conditioned place preference, the time spent on the CS+ floor during the test and the habituation/pre-test sessions (i.e. time-shift) within each treatment group, were analysed with a paired permutation t-test based on 5000 reshuffles of the data, and the data on AMPA:NMDA ratios were additionally analysed with unpaired permutation t-tests against the shared control saline, also based on 5000 reshuffles of the data. The secondary analyses are included in the figures, and the statistical result descriptions can be found on the Supplementary Material. More detailed statistical results of all the tests are available on the statistical tables on the Supplementary Material.

## 3 RESULTS

### 3.1 Mixed evidence of induction of conditioned place preference by LSD and 25CN-NBOH

#### 3.1.1 Unbiased designs

In the initial experiment, CPP1, saline-treated mice showed no clear preference to either side measured as the average time spent on the grid floor (s/min; Figure 1A) with mean difference of 0.3 s/min [CI95% -4.0, 4.6] between the two Floor subgroups. There was a slight tendency in the LSD group mice to spend more time on the drug-paired floor (mean difference 3.6 s/min [CI95% -0.7, 7.9]) and the difference between the subgroups was even larger in the 25CN-NBOH-treated mice (mean difference 8.3 s/min [CI95% 4.0, 12.6]). The two-way ANOVA revealed a significant Floor × Treatment interaction (F(2,24)=3.68, p=0.04), but no significant main effect for the Treatment (F(2,24)=0.55, p=0.6). Further comparisons revealed a significant difference between the 25CN-NBOH-treated subgroups (Holm corrected p=0.009), but not between the saline (Holm corrected p=1.0) nor between the LSD-treated subgroups (Holm corrected p=0.9).

**Figure 1.**
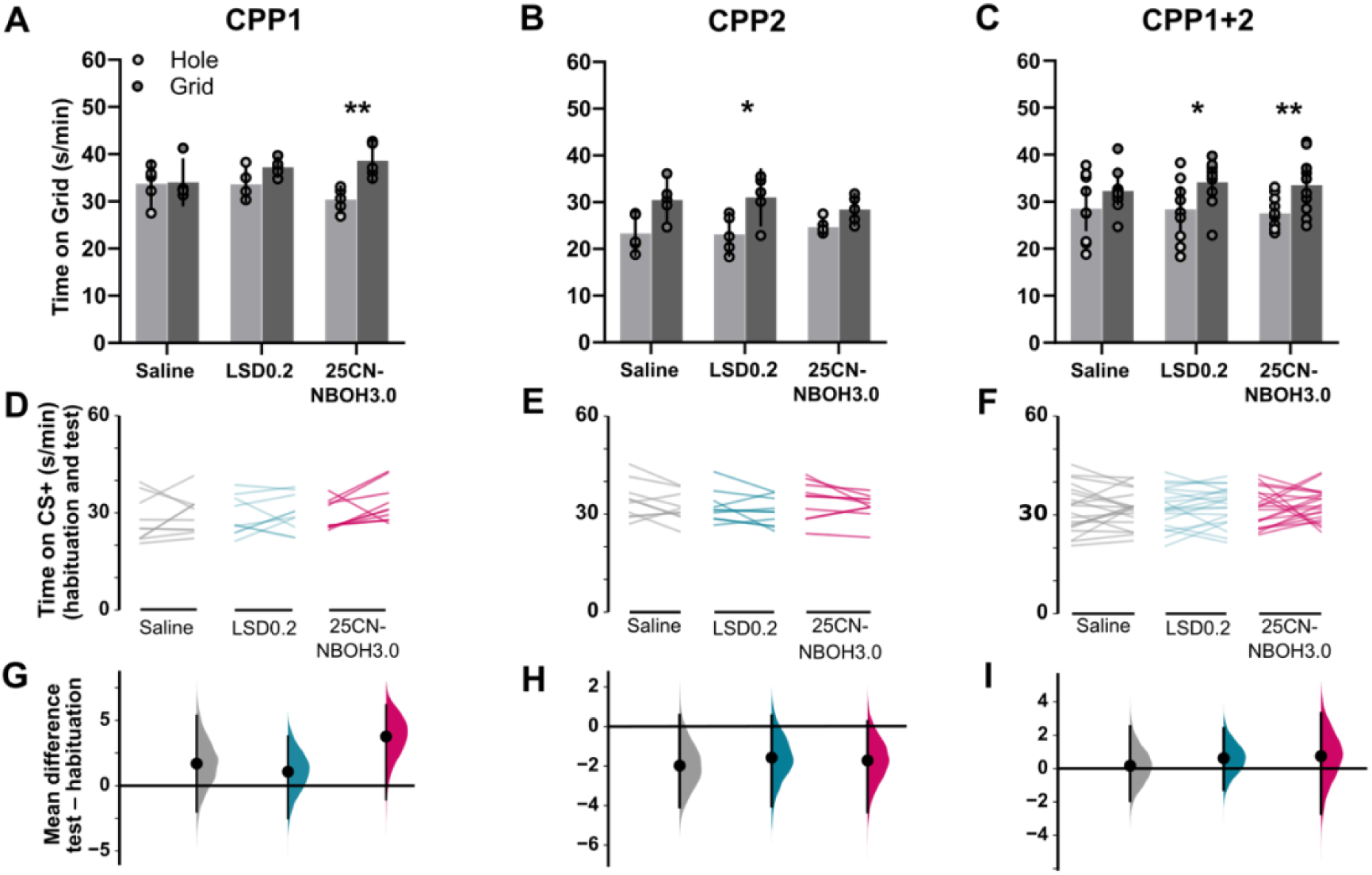
The initial conditioned place preference experiment CPP1 indicated that 3.0 mg/kg 25CN-NBOH would induce conditioned place preference (A) as measured by the within-treatment Floor subgroup difference in the time spent on the Grid. This was also reflected to an extent by increases, although statistically non-significant, in the time spent on the drug-paired side of the apparatus before and after the conditioning (D, G). Replication of the experiment, CPP2, failed to show the same effect; there all the treatments showed a difference between the Floor subgroups with statistically significant effect in 0.2 mg/kg LSD treatment group (B). However, there were no observable changes in the time spent on the drug-paired side in any of the treatment groups (E, F). Pooling the results together maintained the statistically significant differences in the Floor subgroup analysis with both LSD and 25CN-NBOH (C), but the time-shift analysis showed more neutral results (F, I). Data shown as means (bars in A–C; black circles G–I) with 95% confidence intervals (black error bars A–C and G–I) or as individual values (circles A–C; lines D–F). In G–I the paired mean difference is plotted with a bootstrap sampling distribution. * p<0.05, ** p<0.01 difference between the time on Grid between the Grid and Hole subgroups within a treatment group.

In CPP2, the replication of the first experiment, subgroup differences were observed in all treatment groups, saline (mean difference 7.2 s/min [CI95% 2.3, 12.1]), LSD (mean difference 7.9 s/min [CI95% 3.0, 12,8]), and 25CN-NBOH (mean difference 3.7 s/min [CI95% -1.2, 8,6]), as shown in Figure 1B. The two-way ANOVA did not reveal a significant main effect for the Treatment (F(2,24)=0.05, p=1.0) nor Floor × Treatment interaction (F(2,24)=0.9, p=0.4). However, the follow-up multiple comparison showed a significant subgroup difference in the LSD-treated mice (Holm corrected p=0.04), but not in the 25CN-NBOH-(Holm corrected p=0.9) or saline-treated mice (Holm corrected p=0.07).

Pooling the data from these two experiments together, depicted in Figure 1C, still maintained the differences between the conditioning subgroups in the saline-treated mice (mean difference 3.7 s/min [CI95% 0.4, 7.0]), as well as in LSD-(mean difference 5.7 s/min [CI95% 2.4, 9.0]) and 25CN-NBOH-treated mice (mean difference 6.0 s/min [CI95% 2.7, 9.3]). The statistical analysis with two-way ANOVA showed no significant main effect for the Treatment (F(2,53)=0.3, p=0.7) nor for the Floor × Treatment interaction (F(2,53)=0.6, p=0.6). The main effect of the blocking variable Experiment was revealed to be highly significant (F(1,53)=66.6, p<0.0001). The pairwise comparisons further revealed significant differences between the LSD-treated (Holm corrected p=0.01) and the 25CN-NBOH-treated subgroups (Holm corrected p=0.008), but not between the saline-treated subgroups (Holm corrected p=0.19).

In CPP3, using two different doses of LSD, the difference between the Floor subgroups was increased in the lower-dose LSD-treated mice (0.1 mg/kg; mean difference 2.8 s/min [CI95% -5.9, 11.5], Figure 2A), but not in the higher-dose LSD group (0.2 mg/kg; 0.85 s/min [CI95% -7.9, 9.6]) or in the saline control group (mean difference -0.7 s/min [CI95% -9.5, 78.0]). The analysis with the two-way ANOVA revealed no significant main effect for the Treatment (F(2,18)=0.03, p=1.0). The Holm corrected pairwise comparisons did not show any statistical differences within the Treatment groups either (all p=1.0). The difference between the subgroups was increased in the morphine-treated group (mean difference 3.8 s/min [CI95% -4.9, 12.5]), but the statistical tests showed no significant main effects (Treatment F(1,12)=0.3, p=0.6) nor for the pairwise comparisons (all p=1.0).

**Figure 2.**
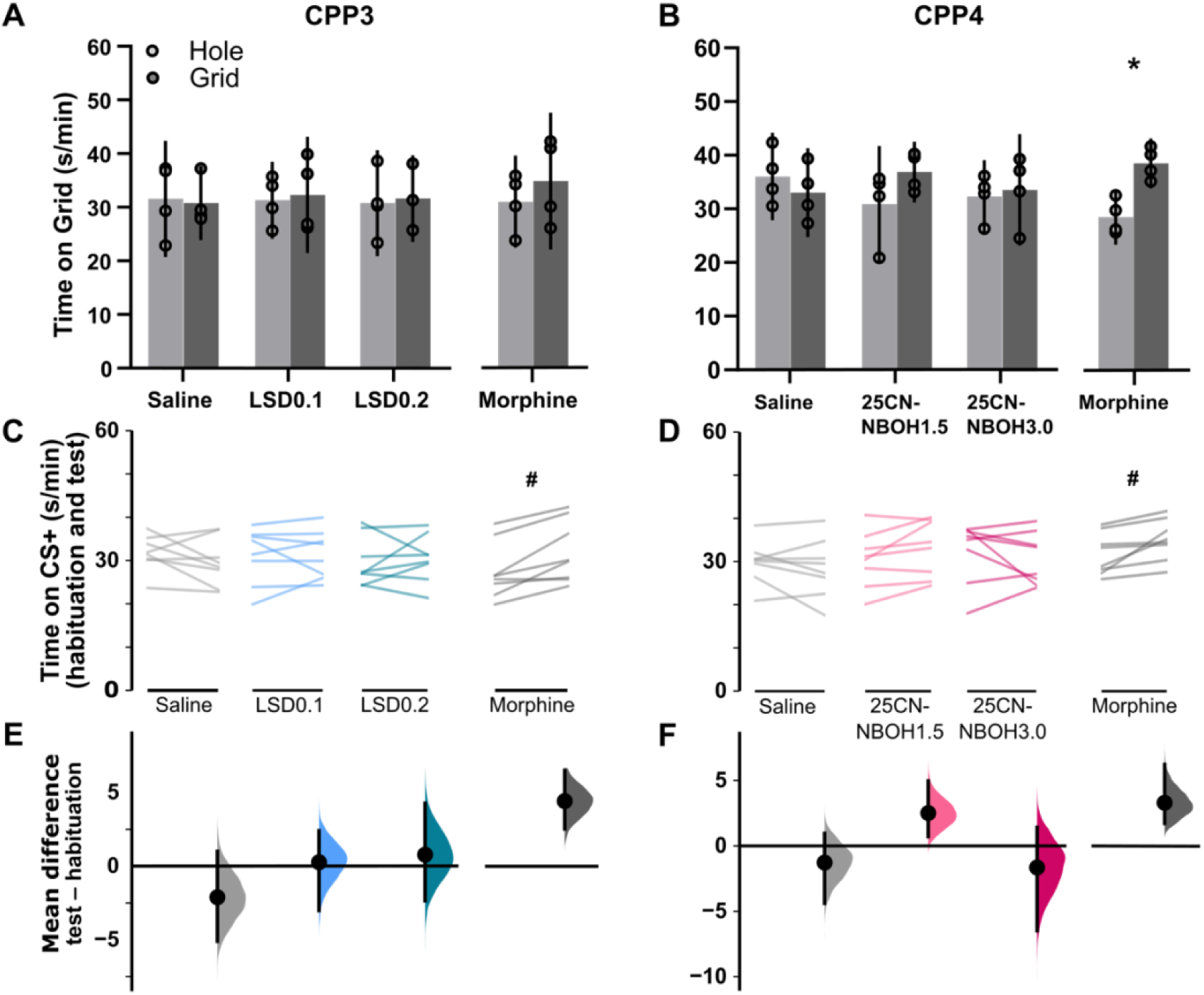
Further analysis of the conditioned place preference with additional doses and morphine as a positive control (CPP3 panels ACE, and CPP4, panels BDF) showed no statistically significant differences between the Floor subgroups with any of the used doses of the serotonergic agents (A, B). Analysing the change in the time spent on the drug-paired side revealed statistically significant increases in the morphine-treated groups in both experiments (C, D), but not with the other treatments. The 1.5 mg/kg 25CN-NBOH showed similar increases in the time spent on the drug-paired floor as morphine (D, F) but the difference did not reach the set significance level. Data shown as means (bars in A–B; black circles E–F) with 95% confidence intervals (black error bars A–B and E–F) or as individual values (circles A–C; lines C–D). In E and F, the paired mean difference is plotted with a bootstrap sampling distribution. * p<0.05 difference between the Grid and Hole subgroups within a treatment group. # p<0.05 paired permutation t-test between the time on drug-paired side on the habituation and the test sessions.

The identical experiment with 25CN-NBOH (CPP4; Figure 2B) showed greater differences between the Floor subgroups in the morphine-treated mice (mean difference 9.9 s/min [CI95% 2.8, 17.14]) and with the lower 25CN-NBOH-treated group (1.5 mg/kg; mean difference 6.0 s/min [CI95% -1.2, 13.1]) while the differences in the saline-treated (mean difference -3.0 [CI95% -10.2, 4.2]) and the group with the higher dose of 25CN-NBOH (3.0 mg/kg; mean difference 1.2 [CI95% - 5.9, 8.4]) were smaller. For the 25CN-NBOH, the two-way ANOVA revealed no significant main effect by the Treatment (F(2,18)=0.2, p=0.8) nor a significant Floor × Treatment interaction (F(2,18)=1.4, p=0.3). The further Holm-corrected pairwise comparisons within the treatment groups were also all non-significant (p=1.0). Analysis for the positive control showed a significant Floor × Treatment interaction (F(1,12)=9.2, p=0.01) and a significant difference between the floor subgroups within the morphine-treated group (Holm corrected p=0.04).

#### 3.1.2 Biased designs

The experiment studying LSD with the biased design, CPP5, showed all the treatments to cause positive time-shifts (saline, 4.7 s/min [CI95% 2.8, 6.3]; morphine, s/min [CI95% 4.6, 10.1]; LSD 0.1 mg/kg, 3.8 s/min [CI95% 1.0, 7.0]; LSD 0.2 mg/kg, 5.2 s/min [CI95% 1.7, 7.5]; Figure 3AE). The statistical analysis of the LSD treatment groups with one-way ANOVA did not reveal any significant effect for the Treatment (F(2,20)=0.32, p=0.7). None of the follow-up pairwise comparisons against the saline control group showed statistically significant differences either (all Holm corrected p≥0.7). Same was the case with the analysis on morphine (t(11.8)=-1.3, p=0.21).

**Figure 3.**
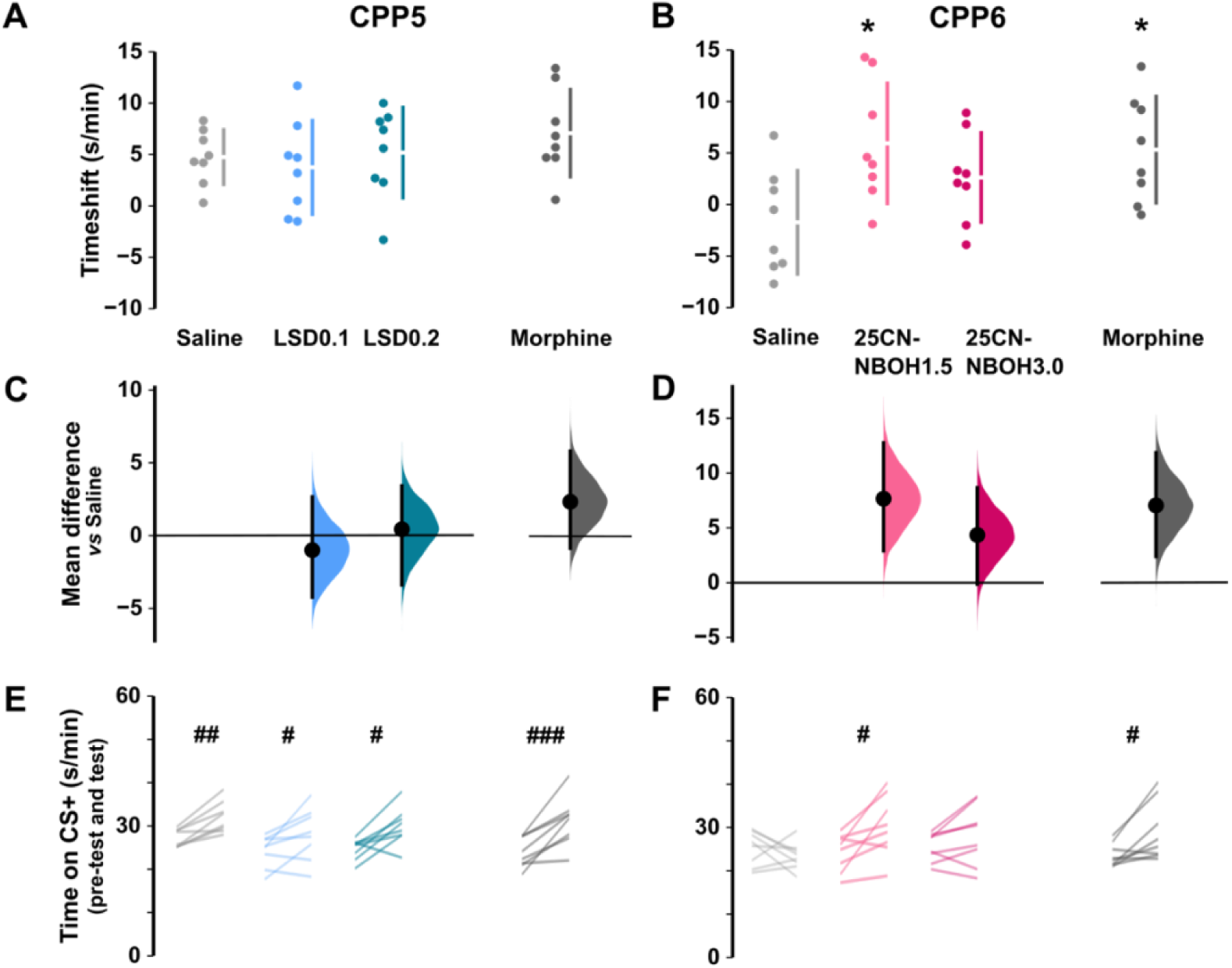
The biased conditioned placed preference experiments (CPP5 panels ACE, and CPP6 panels BDE) showed LSD to cause mild positive time-shifts (A) and slightly more pronounced in the 25CN-NBOH–treated groups (B). From the serotonin agonists, the 1.5 mg/kg 25CN-NBOH caused a statistically significant increase with a time-shift on the similar level as morphine (B, D). In the CPP5, all the treatment groups tended to increase their time spent on the drug-paired floor (E), whereas this was observable only with the 1.5 mg/kg dose of 25CN-NBOH and the positive control morphine in CPP6 (F). Data shown as means (the gap between the bars A–B; black circles C–D) with 95% confidence intervals (bars A–D) or as individual values (dots A–B; lines E–F). In C and D, the unpaired mean difference against saline is plotted with a bootstrap sampling distribution. * p<0.05 Holm corrected pairwise comparison against the saline group time-shift; # p<0.05, ## p<0.01 ### p<0.0001 paired permutation t-test between the time on drug-paired floor on the pre-test and the test sessions.

In CPP6, both morphine (mean time-shift 5.3 s/min [CI95% 1.0, 9.6]) and the lower dose of 25CN-NBOH (1.5 mg/kg; 5.9 s/min [CI95% 1.1, 10.8]) showed clearly positive time-shifts, whereas the changes with the higher dose of 25CN-NBOH (3.0 mg/kg; 2.6 s/min [CI95% -1.0, 6.3]) were less pronounced and in the saline control group (-1.7 s/min [CI95% -5.9, 2.5]) slightly negative, as depicted in Figure 3BF. The one-way ANOVA on the effects of 25CN-NBOH revealed a significant main effect for the Treatment (F(2,20)=4.48, p=0.02), and the post-hoc pairwise comparisons against the saline group showed statistically significant differences for the 1.5 mg/kg 25CN-NBOH (Holm corrected p=0.01) treatment group. Positive control analysis of the morphine-treated group against the saline control group also revealed a statistically significant difference (t(14)=-2.8, p=0.02).

### 3.2 Neither LSD nor 25CN-NBOH caused psychomotor stimulation

Psychomotor stimulation caused by the conditioning interventions in CPP5 and CPP6 was assessed by analysing the total distance moved during each of the 30-min conditioning sessions. These analyses revealed increased locomotor activity in the morphine-treated groups during the CS+ conditioning trials both in CPP5 (Figure 4; Treatment main effect F(1,13)=192, p<0.0001; Holm corrected multiple comparisons morphine *vs.* saline p<0.001) and in CPP6 (Treatment main effect F(1,13)=32.5, p>0.0001; Holm corrected multiple comparisons morphine *vs.* saline p<0.001), while activity in the LSD and 25CN-NBOH groups seemed to stay at the level of the saline control group (CPP5 Treatment main effect F(2,20)=0.6, p=0.6, CPP6 Treatment main effect F(2, 20)=3.6, p=0.05, all pairwise comparisons p>0.07). There were no differences between the treatment groups during the CS– conditioning sessions (Treatment main effect: CPP5 F(3,27)=0.71, p=0.56; CPP6 F(3,27)=2.21, p=0.11).

**Figure 4.**
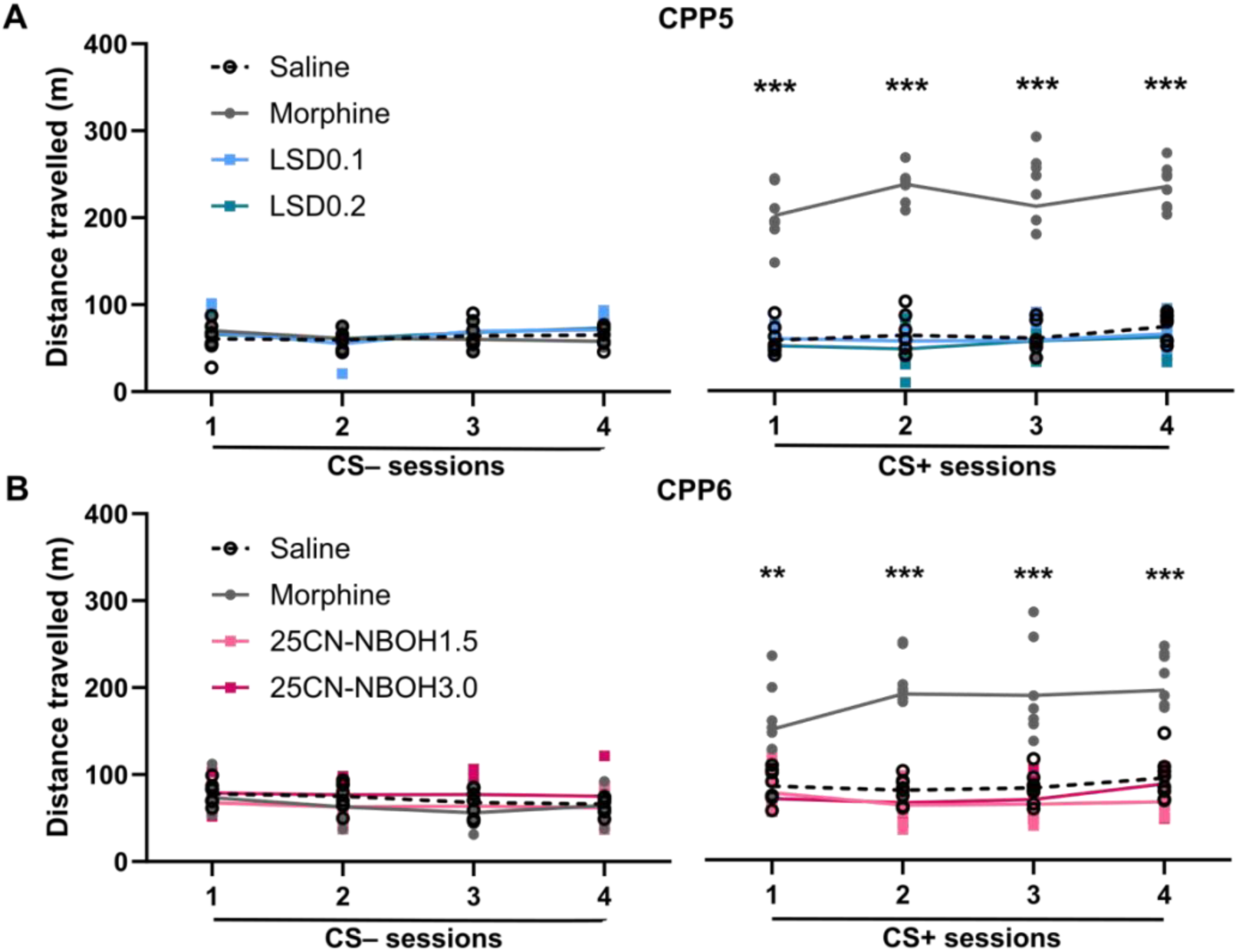
The drug-induced locomotor activation was assessed from the conditioning sessions of the CPP5 (A) and CPP6 (B). In both experiments, the morphine treatment caused significant increases in the locomotor activity during the CS+ sessions but with all investigated doses of LSD and 25CN-NBOH the activity levels were similar as those in the CS– sessions with saline injections. The data is shown as means (lines) with individual values (dots). ** p<0.01, *** p<0.001 Holm corrected pairwise comparison against saline group mean.

### 3.3 LSD, but not 25CN-NBOH, induced glutamatergic plasticity in dopamine neurons in the VTA

For the electrophysiological measurements, all the comparisons were made against the AMPA:NMDA ratio of the saline-treated mice (average mean ratio 0.28 [CI95% 0.19, 0.37], n=15; lateral population mean 0.29 [CI95% 0.18, 0.41], n=13; medial population mean 0.29 [CI95% 0.07, 0.52], n=14).

When analysing the average ratios, LSD treatment induced clearly increased AMPA:NMDA ratios (mean ratio 0.7 [CI95% 0.32, 1.08], n=11; Figure 5AB) in comparison to the saline-treated group, whereas the ratios in the 25CN-NBOH group were on the same level as the saline group’s (mean ratio 0.40 [CI95% 0.2, 0.6], n=7). The one-way ANOVA revealed a significant Treatment main effect (F(2,29)=6.1, p=0.006) and the further pairwise comparisons revealed the difference between the saline and LSD treatment groups to be statistically significant (mean difference 0.4, Holm corrected p=0.003) whereas the increment in 25CN-NBOH treated group was not (mean difference 0.1, Holm corrected p=0.18). Also, morphine significantly increased the AMPA:NMDA ratio (morphine mean ratio 0.57 [CI95% 0.05, 1.1], n=4; mean difference 0.3, t(17)=-2.5, p=0.02).

**Figure 5.**
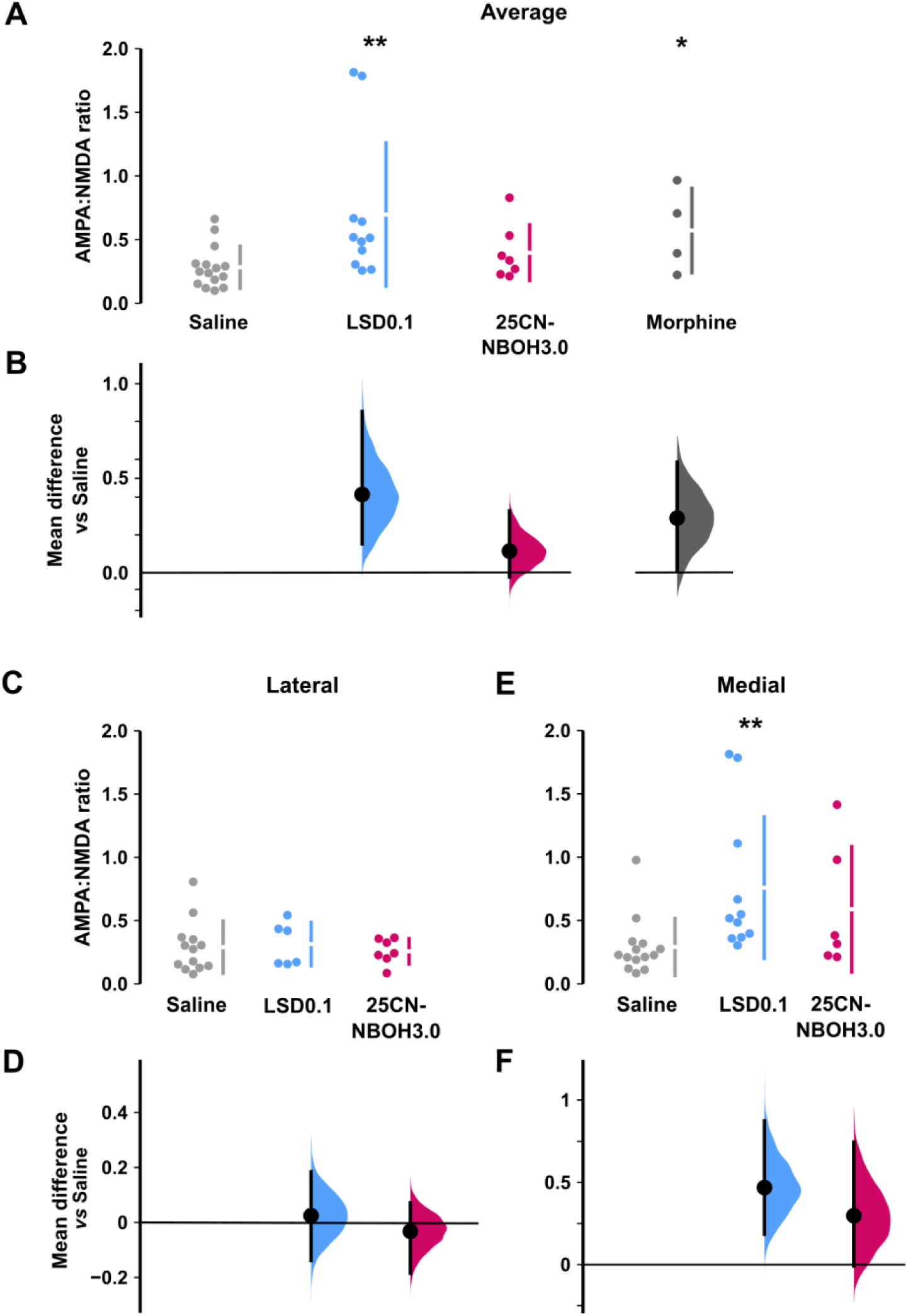
The ratio between the AMPA and NMDA receptor currents in the VTA dopamine neurons assessed with whole-cell patch-clamp electrophysiology. Comparing the average ratios (1-2 cells per animal; AB), treatment with 0.1 mg/kg LSD caused a clear and significant increase, on par with the positive control morphine. The effects seemed to emerge from an anatomically distinct population since in the lateral population of neurons, neither of the serotonergic agents induced any observable changes in the ratio (CD). However, in the medial population, 0.1 mg/kg LSD again showed significant increases in the AMPA:NMDA ratio (EF), and while there was an increase also with 3.0 mg/kg 25CN-NBOH, the difference did not reach the set level of significance. The data shown as means (the gap between the bars in ACE; black circles BDF) with 95% confidence intervals (bars A–F) and as individual values (dots ACE). In B, D and F the unpaired mean difference against saline is plotted with a bootstrap sampling distribution. * p<0.05 ** p<0.01 pairwise comparison against the saline group.

In the lateral dopamine neuron population, neither LSD (0.31 [CI95% 0.15, 0.48], n=6) or 25CN-NBOH (0.26 [CI95% 0.12, 0.40], n=7) showed any great difference compared to the saline control (Figure 5CD). The statistical analysis with one-way ANOVA did not reveal statistically significant main effect for Treatment (F=2,22)=0.14, p=0.9), and further pairwise comparisons against the saline group showed no differences between the treatment groups that would have reached the specified significance level (all Holm corrected p=1.0).

In the medial VTA neurons, treatment with both LSD and 25CN-NBOH induced increased AMPA:NMDA ratios, LSD treatment showing clearer increases (0.76 [CI95% 0.51, 1.01], n=11) while that of 25CN-NBOH remaining more modest (0.59 [CI95% 0.31, 0.87], n=6; Figure 5EF). One-way ANOVA showed a statistically significant Treatment main effect (F(2, 27)=6.5, p=0.005), and Holm’s corrected pairwise comparison revealed significant differences between the saline and LSD groups (p=0.003), but not between the saline and 25CN-NBOH groups (p=0.07).

## 4 DISCUSSION

In this study attempting to understand their potential rewarding properties in mouse models, we found mixed results of the classical psychedelic LSD and the selective 5-HT_2A_ receptor agonist 25CN-NBOH in inducing CPP and glutamatergic synaptic plasticity in dopamine neurons of the VTA. The main outcomes of the experiments are summarised in Table 2.

**Table 2.** An overview of the findings on the effects of the serotonergic agonists in each experiment. For the CPP, the primary and secondary criteria are described in Table 1 and the statistical results descriptions can be found on Supplementary Material.

| **Experiment** | **Evidence of conditioned place preference** |  |
| --- | --- | --- |
|  | **Primary** | **Secondary** |
| CPP1 | 3.0 25CN-NBOH | No |
| CPP2 | 0.2 LSD | No |
| CPP1+2 pooled | 0.2 LSD, 3.0 25CN-NBOH | No |
| CPP3 | No | No |
| CPP4 | No | No |
| CPP5 | No | All |
| CPP6 | 1.5 25CN-NBOH | 1.5 25CN-NBOH |
| **Locomotor activity** | **Evidence of psychomotor stimulation** |  |
| LSD (CPP5) | No |  |
| 25CN-NBOH (CPP6) | No |  |
| **AMPA:NMDA ratio** | **Evidence of increase** |  |
| Average | 0.1 LSD |  |
| Lateral population | No |  |
| Medial population | 0.1 LSD |  |

Our initial experiment CPP1 was originally ran as a control experiment for another study with an aim to show that LSD and 25CN-NBOH do not induce CPP in mice, yet the tested 3.0 mg/kg dose of 25CN-NBOH induced place preference (Figure 1A), initiating the rest of the study. However, this effect with the 3.0 mg/kg 25CN-NBOH was not repeated in any of our following experiments, whereas the lower dose of 1.5 mg/kg of the drug did show either significant preference formation (CPP6, Figure 3BDF) or tendencies towards it (CPP4, Figure 2DF). LSD, on the other hand, showed any form of preference induction only in CPP2 with the dose of 0.2 mg/kg (Figure1B), but not even tendencies in the later experiments. These data on LSD seem to contradict the main claims of the earlier publications by Parker (1996) and Meehan and Schechter (1998), both of which report that LSD produces conditioned place preference in rats. At a closer look, these results also had several inconsistencies: in Parker’s study only the 0.2 mg/kg dose of LSD produced place preference whereas the lower doses (0.025–0.1 mg/kg) failed to do so. Intriguingly, the preference was produced only when the tests were run without any pre-exposure to the conditioning arena and introducing a 15-min pre-test session prevented the induction of place preference. This effect where an exposure to the CS in the absence of the US can delay or completely abolish the conditioning is called latent inhibition (Tzschentke, 1998) and it could explain the lack of consistent CPP expression in our study as all our CPP experiments included habituation or pre-test sessions. However, Meehan and Schechter used a design where the animals had two 15-min sessions of free access to the conditioning apparatus before the conditioning phase and the 0.2 mg/kg dose of LSD still produced CPP, indicating that latent inhibition effect is not as strong in inhibiting the preference induction as Parker’s findings initially implied. In the Meehan and Schechter study the preference was also only observed in male and not in female rats. In our data, there was no implication of any sex differences in any of the responses, although it is worth noting that we used the sex only as a blocking factor and the experiments were not powered to observe other than the most drastic sex differences. Based on our results, we cannot conclude that LSD or 25CN-NBOH would never produce CPP in mice and there certainly might be some circumstances and conditions where the drugs have rewarding properties strong enough to produce conditioned approach behaviour. However, based on these results, neither of the tested drugs seem to be so highly positively reinforcing that they would induce place preference reliably, as compared to morphine. This verdict echoes the one of Parker who concluded that “the strength of a place preference produced by LSD appears to be weak relative to other rewarding drugs” (Parker, 1996).

The time course of the drugs’ effects might also give an interesting perspective on the potentially rewarding effects of the serotonergic agonists. Recently, Vargas-Perez *et al*. (2024) proposed that 4-acetoxy-N,N-dimethyltryptamine (4-AcO-DMT), a prodrug for the psychedelic psilocin (Nichols & Frescas, 1999; Jones *et al*., 2024), would have rewarding stimulus effects both acutely and during the prolonged after-effect period lasting at least until 24 h after the administration. In their study, isolating these two effects with modified experimental designs showed that both the acute and the after-effects of the drug induced place preference in male rats, yet these rewarding effects were not observed when the traditional unbiased and counterbalanced place preference design with 24 h between the drug and saline conditioning sessions was used. The authors proposed that this effect could be due to rewarding nature of the after-effects: if the after-effects, that are present during the CS– session with the vehicle, are as rewarding or even more rewarding than the acute effects present at the CS+ sessions, the animals might tend to approach the vehicle-paired environment where these after-effects originally took place. In a normal, unbiased and counter-balanced CPP design this would mean that the animals have been conditioned to associate rewarding drug stimuli with both conditioning environments which would also lead to approach behaviour towards both of them when tested. This would then cause a lack of clearly observable preference, and thus end up looking as if no drug reward association was formed during the conditioning. If we extrapolate these findings to the present study, postulating that the effects were to be true with LSD and 25CN-NBOH and in mice, such rewarding after-effects would have been present in the unbiased experiments (CPP1-4) during either all or half of the saline conditioning sessions and potentially during the test session depending on the counterbalanced schedule the mouse was part of. In the biased designs (CPP5-6) the same holds true if we assume that the after-effects are present already before the 24-h measuring point. These effects, if present, could have affected the conditioning and the formation of the place preference. The findings by Vargas-Perez et al. might also reflect interestingly the findings of Marona-Lewicka *et al*. (2005, 2009) who showed that the discriminatory stimulus effects of LSD are initially serotonin-dependent, but modulated more by the dopamine system at the later time-points. Both the existence of the reinforcing after-effects, their relation to the dopamine system activation, and to the formation of place preference with LSD and other psychedelics warrant further experiments to demonstrate.

The drugs’ effects on the locomotor activity of the mice were also measured in the two last CPP experiments. While morphine prominently increased the locomotor activity during the conditioning sessions, no effects were observed with any of the tested doses of LSD and 25CN-NBOH. Additionally, the data also showed no evidence of either sensitisation or tolerance formation during the week-long repeated administration. Locomotor activity is commonly measured together with the main outcome measures in CPP experiments as an important control measure (Cunningham *et al*., 2006), but many drugs of abuse, such as stimulants, nicotine, and to an extent ethanol, in addition to opioids (morphine used here), are known to induce both psychomotor stimulation and CPP (Tzschentke, 1998). There is also a known connection between the increases in striatal dopamine release and increases in locomotor activity, originally a basis for the (now largely refuted) psychomotor stimulant theory of addiction (Wise & Bozarth, 1987). Following this logic, neither of the serotonergic agonists tested here would be positively reinforcing, echoing the main outcomes of the CPP studies. However, changes in locomotor activity are known not to be a good predictor for a drug’s ability to cause CPP and the rewarding effects of several drugs have been shown to be separable from both the locomotor effects and the striatal dopamine actions (Pettit *et al*., 1984; Vaccarino *et al*., 1986; Carr *et al*., 1988; Risinger *et al*., 1992; Tzschentke, 1998). Therefore, any conclusions about the rewarding effects of a drug based on the locomotor activity should be made with caution.

Some limitations should be acknowledged when interpreting the present CPP data. First, even if both sexes were tested in the biased design CPP and the electrophysiological experiments, most of the tests were conducted only in male mice. This consolidates the well-acknowledged bias in the scientific literature (Shansky & Murphy, 2021), limits the generalisability of the present findings and warrants further studies to properly include sex as a biological variable. Second, there seemed to be a fairly strong effect of either the investigator or the time of the experiment on the behavioural responses, most evident between the experiments CPP1 and CPP2: the apparatus, the strain of mice and their vendor, as well as the design of the study were all identical, but the experiment was conducted by a different investigator, which evidently caused some bias in the output (shown by the tendency for a difference in the time on Grid in the negative control saline in Figure 1B, even if not statistically significant). This is supported by the significant effect of the blocking factor ‘Experiment’ used in the analysis of the pooled data from CPP1 and CPP2 (see Results 3.1.1.). Stressors and additional stimuli are known to affect the main outcome measures of the test and the formation of associations (Tzschentke, 1998; although see Bechtholt *et al*., 2004). While Cunningham’s work has shown that using counterbalanced, unbiased study design should yield reliable results even if the study population has a bias on the used setup (Cunningham *et al*., 2003, 2006), the secondary outcome measures were added to the analysis to give a more comprehensive perspective on the data. Despite these limitations, we consider that our behavioural data do not support clear reinforcement by acute repeated dosing of psychedelics and 5HT_2A_ receptor agonists in mice.

Many drugs of abuse known to be positively reinforcing have been shown to modify synaptic plasticity, commonly emerging as increased ratio between AMPA and NMDA receptor-mediated excitatory currents, in the dopamine releasing neurons of the VTA, when studied *ex vivo* one or several days after a single drug dose (Lüscher & Malenka, 2011). Previous studies also suggest an anatomical segregation of dopamine neurons: an antero-lateral population that reacts exclusively to rewarding stimuli, and a medio-posterior population that responds to both aversive and rewarding stimuli, presumably responsible for detecting salience in general (Lammel *et al*., 2011). Our data showed that a single 0.1 mg/kg dose of LSD, but not the 3.0 mg/kg dose of 25CN-NBOH, induced moderate plasticity in the VTA dopamine neurons. Anatomically divided analysis further showed that the LSD treatment caused significant increases in the medially situated dopamine neurons while no changes were observed in the lateral populations. Even if not statistically significant, also the treatment with 25CN-NBOH approximately doubled the AMPA:NMDA ratio compared to that of saline control. There is a strong correlation between the capability of a drug to induce increased AMPA:NMDA ratios and cause CPP as these synaptic adaptations have been shown to be triggered by psychostimulants, morphine, nicotine, ethanol, and benzodiazepines (Saal *et al*., 2003; Heikkinen *et al*., 2009; Vashchinkina *et al*., 2018), all of which have also been shown to produce place preference (reviewed broadly in Tzschentke, 1998). Similarly, interfering with the glutamatergic plasticity in the VTA has been shown to hamper the induction of place preference (Harris & Aston-Jones, 2003; Harris *et al*., 2004). On the other hand, there are several examples showing that physical stress, formalin irritation, and drugs producing conditioned place aversion instead of preference also generate increases in AMPA:NMDA ratios on VTA dopamine neurons, predominantly in the medio-posterior dopaminergic population (Saal *et al*., 2003; Lammel *et al*., 2011; Vashchinkina *et al*., 2012, 2014). Furthermore, the medial dopamine neurons have been shown to send axonal projections at least to the basolateral amygdala and frontal cortical areas in addition to the striatal targets, and these neurons have projection site specificity in their responses to external stimuli and their valence (Lammel *et al*., 2011). As we lack the projection-specificity of the neurons on which the induction of the glutamatergic plasticity was observed after LSD and 25CN-NBOH, further studies are warranted to fully understand the implication of this effect. Altogether, induction of plasticity on the medial population of VTA dopamine neurons that we observed cannot automatically be linked with rewarding or aversive stimulus properties but could tell about detected salience of the stimuli. Importantly, the finding does not disagree with our behavioural findings: the mixed nature of the valence responsivity especially in the medially situated VTA dopamine neurons is much in line with the mixed results in the CPP experiments.

## 5 CONCLUSIONS

Here we report to our knowledge the first conditioned place preference experiments with classical serotonergic psychedelics in mice. Despite contradicting the earlier findings with LSD in rats, the results with no reliable induction of place preference reflect most earlier literature in indicating either very weak rewarding or mixed rewarding and aversive stimulus properties. This conclusion is supported by the electrophysiological findings showing neuroplastic changes only in glutamatergic inputs to the medial population of the VTA dopamine neurons, known to also have mixed responsivity to rewarding and aversive stimuli, being different from robust plasticity produced by most positively reinforcing drugs of abuse, such as opioids and stimulants.

## Acknowledgements

We want to thank Heidi Hytönen and Annika Schäfer for their assistance, and Dr. Jaakko Kopra, Dr. Petteri Piepponen, and Prof. Mikko Airavaara for the access to and guidance with the place preference apparatus.

## Author Disclosures

### Funding information

This work was supported by grants from the Finnish Cultural Foundation (no. 00180226; LVE), the Finnish Foundation for Alcohol Studies (LVE, ERK), Instrumentarium Science Foundation (LVE), Paulo Foundation (LVE), and Research Council of Finland (no.1317399; ERK). None of the funders had any role in the design of the study, in the collection, analysis or the interpretation of the data, writing of the report nor in the decision to submit the paper for publication.

### CRediT author contributions

Conceptualisation: LVE, EN, ERK

Funding acquisition: LVE, ERK

Supervision: LVE, ERK

Investigation: LVE, J-PL (behavioural), EN (electrophysiological)

Formal analysis: LVE

Visualisation: LVE

Writing – original draft: LVE

Writing – review and editing: LVE, EN, J-PL, ERK

### Conflict of Interest

The authors declare that they have no conflicts of interest.

### Data availability

The data that support the findings of this study are openly available in Open Science Framework (OSF) repository at http://doi.org/10.17605/OSF.IO/4QRKC.

## Notes

http://doi.org/10.17605/OSF.IO/4QRKC

### Competing Interest Statement

The authors have declared no competing interest.

## REFERENCES

Aitta-aho, T., Möykkynen, T.P., Panhelainen, A.E., Vekovischeva, O.Y., Bäckström, P., & Korpi, E.R. (2012) Importance of GluA1 Subunit-Containing AMPA Glutamate Receptors for Morphine State-Dependency. PLOS ONE, 7, e38325.

Amsterdam, J. van, Opperhuizen, A., & Brink, W. van den (2011) Harm potential of magic mushroom use: A review. Regulatory Toxicology and Pharmacology, 59, 423–429.

Baker, L.E. (2018) Hallucinogens in Drug Discrimination. In Halberstadt, A.L., Vollenweider, F.X., & Nichols, D.E. (eds), Behavioral Neurobiology of Psychedelic Drugs. Springer, Berlin, Heidelberg, pp. 201–219.

Basedow, L.A. & Kuitunen-Paul, S. (2022) Motives for the use of serotonergic psychedelics: A systematic review. Drug and Alcohol Review, 41, 1391–1403.

Bate, S.T. & Clark, R.A. (2014) The Design and Statistical Analysis of Animal Experiments, 1st edn. Cambridge University Press.

Bechtholt, A.J., Gremel, C.M., & Cunningham, C.L. (2004) Handling blocks expression of conditioned place aversion but not conditioned place preference produced by ethanol in mice. Pharmacology Biochemistry and Behavior, 79, 739–744.

Berridge, K.C., Robinson, T.E., & Aldridge, J.W. (2009) Dissecting components of reward: ‘liking’, ‘wanting’, and learning. Current Opinion in Pharmacology, 9, 65–73.

Buchborn, T., Lyons, T., & Knöpfel, T. (2018) Tolerance and Tachyphylaxis to Head Twitches Induced by the 5-HT2A Agonist 25CN-NBOH in Mice. Frontiers in Pharmacology, 9, 1– 8.

Carbonaro, T.M., Johnson, M.W., & Griffiths, R.R. (2020) Subjective features of the psilocybin experience that may account for its self-administration by humans: a double-blind comparison of psilocybin and dextromethorphan. Psychopharmacology, 237, 2293–2304.

Carr, G.D., Phillips, A.G., & Fibiger, H.C. (1988) Independence of amphetamine reward from locomotor stimulation demonstrated by conditioned place preference. Psychopharmacology, 94, 221–226.

Chen, B.T., Bowers, M.S., Martin, M., Hopf, F.W., Guillory, A.M., Carelli, R.M., Chou, J.K., & Bonci, A. (2008) Cocaine but Not Natural Reward Self-Administration nor Passive Cocaine Infusion Produces Persistent LTP in the VTA. Neuron, 59, 288–297.

Clark, R.A., Shoaib, M., Hewitt, K.N., Stanford, S.C., & Bate, S.T. (2012) A comparison of InVivoStat with other statistical software packages for analysis of data generated from animal experiments. J Psychopharmacol, 26, 1136–1142.

Cumming, G. (2014) The New Statistics: Why and How. Psychol Sci, 25, 7–29.

Cumming, G. & Calin-Jageman, R. (2017) Introduction to the New Statistics: Estimation, Open Science, and Beyond. Routledge Taylor & Francis Group, London : New York.

Cunningham, C.L., Ferree, N.K., & Howard, M.A. (2003) Apparatus bias and place conditioning with ethanol in mice. Psychopharmacology, 170, 409–422.

Cunningham, C.L., Gremel, C.M., & Groblewski, P.A. (2006) Drug-induced conditioned place preference and aversion in mice. Nat Protoc, 1, 1662–1670.

Cunningham, C.L., Groblewski, P.A., & Voorhees, C.M. (2011) Place Conditioning. In Olmstead, M.C. (ed), Animal Models of Drug Addiction. Humana Press, Totowa, NJ, pp. 167–189.

de Jong, J.W., Afjei, S.A., Pollak Dorocic, I., Peck, J.R., Liu, C., Kim, C.K., Tian, L., Deisseroth, K., & Lammel, S. (2019) A Neural Circuit Mechanism for Encoding Aversive Stimuli in the Mesolimbic Dopamine System. Neuron, 101, 133–151.e7.

Elsilä, L.V., Harkki, J., Enberg, E., Martti, A., Linden, A.-M., & Korpi, E.R. (2022) Effects of acute lysergic acid diethylamide on intermittent ethanol and sucrose drinking and intracranial self-stimulation in C57BL/6 mice. J Psychopharmacol, 36, 860–874.

Elsilä, L.V., Korhonen, N., Hyytiä, P., & Korpi, E.R. (2020) Acute Lysergic Acid Diethylamide Does Not Influence Reward-Driven Decision Making of C57BL/6 Mice in the Iowa Gambling Task. Frontiers in Pharmacology, 11, 1–10.

Fantegrossi, W.E., Gray, B.W., Bailey, J.M., Smith, D.A., Hansen, M., & Kristensen, J.L. (2015) Hallucinogen-like effects of 2-([2-(4-cyano-2,5-dimethoxyphenyl) ethylamino]methyl)phenol (25CN-NBOH), a novel N-benzylphenethylamine with 100-fold selectivity for 5-HT2Areceptors, in mice. Psychopharmacology, 232, 1039– 1047.

Fantegrossi, W.E., Woods, J.H., & Winger, G. (2004) Transient reinforcing effects of phenylisopropylamine and indolealkylamine hallucinogens in rhesus monkeys: Behavioural Pharmacology, 15, 149–157.

Franklin, K.B.J. & Paxinos, G. (2008) The Mouse Brain in Stereotaxic Coordinates, 3rd edn. Academic Press.

Glynos, N.G., Fields, C.W., Barron, J., Herberholz, M., Kruger, D.J., & Boehnke, K.F. (2023) Naturalistic Psychedelic Use: A World Apart from Clinical Care. Journal of Psychoactive Drugs, 55, 379–388.

Gong, S., Zheng, C., Doughty, M.L., Losos, K., Didkovsky, N., Schambra, U.B., Nowak, N.J., Joyner, A., Leblanc, G., Hatten, M.E., & Heintz, N. (2003) A gene expression atlas of the central nervous system based on bacterial artificial chromosomes. Nature, 425, 917–925.

Goodwin, A.K. (2016) An intravenous self-administration procedure for assessing the reinforcing effects of hallucinogens in nonhuman primates. Journal of Pharmacological and Toxicological Methods, 82, 31–36.

Gouveia, K. & Hurst, J.L. (2017) Optimising reliability of mouse performance in behavioural testing: the major role of non-aversive handling. Sci Rep, 7, 44999.

Grieco, S.F., Castrén, E., Knudsen, G.M., Kwan, A.C., Olson, D.E., Zuo, Y., Holmes, T.C., & Xu, X. (2022) Psychedelics and Neural Plasticity: Therapeutic Implications. Journal of Neuroscience, 42, 11.

Griffiths, R.R., Brady, J.V., & Bradford, L.D. (1979) Predicting the Abuse Liability of Drugs with Animal Drug Self-Administration Procedures: Psychomotor Stimulants and Hallucinogens. In Thompson, T. & Dews, P.B. (eds), Advances in Behavioral Pharmacology. Elsevier, pp. 163–208.

Halberstadt, A.L., Chatha, M., Klein, A.K., Wallach, J., & Brandt, S.D. (2020) Correlation between the potency of hallucinogens in the mouse head-twitch response assay and their behavioral and subjective effects in other species. Neuropharmacology, 167, 107933.

Halberstadt, A.L. & Geyer, M.A. (2013) Characterization of the head-twitch response induced by hallucinogens in mice: Detection of the behavior based on the dynamics of head movement. Psychopharmacology, 227, 727–739.

Hall, G. (1994) Pavlovian Conditioning. In Mackintosh, N.J. (ed), Animal Learning and Cognition, 2nd edn, Handbook of Perception and Cognition. Academic Press, San Diego, CA, USA.

Harris, G.C. & Aston-Jones, G. (2003) Critical Role for Ventral Tegmental Glutamate in Preference for a Cocaine-Conditioned Environment. Neuropsychopharmacol, 28, 73– 76.

Harris, G.C., Wimmer, M., Byrne, R., & Aston-Jones, G. (2004) Glutamate-associated plasticity in the ventral tegmental area is necessary for conditioning environmental stimuli with morphine. Neuroscience, 129, 841–847.

Heikkinen, A.E., Möykkynen, T.P., & Korpi, E.R. (2009) Long-lasting Modulation of Glutamatergic Transmission in VTA Dopamine Neurons after a Single Dose of Benzodiazepine Agonists. Neuropsychopharmacol, 34, 290–298.

Ho, J., Tumkaya, T., Aryal, S., Choi, H., & Claridge-Chang, A. (2019) Moving beyond P values: data analysis with estimation graphics. Nat Methods, 16, 565–566.

Hoffmeister, F. (1975) Negative reinforcing properties of some psychotropic drugs in drug-naive rhesus monkeys. J Pharmacol Exp Ther, 192, 468–477.

Holze, F., Ley, L., Müller, F., Becker, A.M., Straumann, I., Vizeli, P., Kuehne, S.S., Roder, M.A., Duthaler, U., Kolaczynska, K.E., Varghese, N., Eckert, A., & Liechti, M.E. (2022) Direct comparison of the acute effects of lysergic acid diethylamide and psilocybin in a double-blind placebo-controlled study in healthy subjects. Neuropsychopharmacol., 47, 1180–1187.

Hurst, J.L. & West, R.S. (2010) Taming anxiety in laboratory mice. Nat Methods, 7, 825–826.

Jaster, A.M., Elder, H., Marsh, S.A., de la Fuente Revenga, M., Negus, S.S., & González-Maeso, J. (2022) Effects of the 5-HT2A receptor antagonist volinanserin on head-twitch response and intracranial self-stimulation depression induced by different structural classes of psychedelics in rodents. Psychopharmacology,.

Jensen, A.A., Mccorvy, J.D., Leth-Petersen, S., Bundgaard, C., Liebscher, G., Kenakin, T.P., Bräuner-Osborne, H., Kehler, J., & Kristensen, J.L. (2017) Detailed Characterization of the In Vitro Pharmacological and Pharmacokinetic Properties of N-(2-Hydroxybenzyl)-2, 5-Dimethoxy-4-Cyanophenylethylamine (25CN-NBOH), a Highly Selective and Brain-Penetrant 5-HT 2A Receptor Agonist s. The Journal of Pharmacology and Experimental Therapeutics, 361, 441–453.

Johnson, M.W. & Ettinger, R.H. (2000) Active cocaine immunization attenuates the discriminative properties of cocaine. Experimental and Clinical Psychopharmacology, 8, 163–167.

Johnson, M.W., Griffiths, R.R., Hendricks, P.S., & Henningfield, J.E. (2018) The abuse potential of medical psilocybin according to the 8 factors of the Controlled Substances Act. *Neuropharmacology*, Psychedelics: New Doors, Altered Perceptions, 142, 143–166.

Jones, N.T., Wagner, L., Hahn, M.C.P., Scarlett, C.O., & Wenthur, C.J. (2024) In vivo validation of psilacetin as a prodrug yielding modestly lower peripheral psilocin exposure than psilocybin. Front. Psychiatry, 14.

Katsidoni, V., Apazoglou, K., & Panagis, G. (2011) Role of serotonin 5-HT2A and 5-HT2C receptors on brain stimulation reward and the reward-facilitating effect of cocaine. Psychopharmacology, 213, 337–354.

Klawonn, A.M. & Malenka, R.C. (2018) Nucleus Accumbens Modulation in Reward and Aversion. Cold Spring Harb Symp Quant Biol, 83, 119–129.

Konradi, C. & Hurd, Y.L. (2023) Drug Use Disorders and Addiction. In Brunton, L.L. & Knollmann, B.C. (eds), Goodman and Gilman’s The Pharmacological Basis of Therapeutics, 14th edn. McGraw Hill, New York, NY, USA.

Koob, G.F., Arends, M.A., & Le Moal, M. (2014) Chapter 3 - Animal Models of Addiction. In Koob, G.F., Arends, M.A., & Le Moal, M. (eds), Drugs, Addiction, and the Brain. Academic Press, San Diego, pp. 65–91.

Kopra, J., Villarta-Aguilera, M., Savolainen, M., Weingerl, S., Myöhänen, T.T., Rannanpää, S., Salvatore, M.F., Andressoo, J.O., & Piepponen, T.P. (2018) Constitutive Ret signaling leads to long-lasting expression of amphetamine-induced place conditioning via elevation of mesolimbic dopamine. Neuropharmacology, 128, 221–230.

Lammel, S., Ion, D.I., Roeper, J., & Malenka, R.C. (2011) Projection-Specific Modulation of Dopamine Neuron Synapses by Aversive and Rewarding Stimuli. Neuron, 70, 855– 862.

Lammel, S., Lim, B.K., & Malenka, R.C. (2014) Reward and aversion in a heterogeneous midbrain dopamine system. Neuropharmacology, NIDA 40th Anniversary Issue, 76, 351–359.

Ley, L., Holze, F., Arikci, D., Becker, A.M., Straumann, I., Klaiber, A., Coviello, F., Dierbach, S., Thomann, J., Duthaler, U., Luethi, D., Varghese, N., Eckert, A., & Liechti, M.E. (2023) Comparative acute effects of mescaline, lysergic acid diethylamide, and psilocybin in a randomized, double-blind, placebo-controlled cross-over study in healthy participants. Neuropsychopharmacol., 48, 1659–1667.

Lüscher, C. & Malenka, R.C. (2011) Drug-Evoked Synaptic Plasticity in Addiction: From Molecular Changes to Circuit Remodeling. Neuron, 69, 650–663.

Marona-Lewicka, D., Chemel, B.R., & Nichols, D.E. (2009) Dopamine D4 receptor involvement in the discriminative stimulus effects in rats of LSD, but not the phenethylamine hallucinogen DOI. Psychopharmacology, 203, 265–277.

Marona-Lewicka, D., Thisted, R.A., & Nichols, D.E. (2005) Distinct temporal phases in the behavioral pharmacology of LSD: Dopamine D2 receptor-mediated effects in the rat and implications for psychosis. Psychopharmacology, 180, 427–435.

Meehan, S.M. & Schechter, M.D. (1998) LSD produces conditioned place preference in male but not female fawn hooded rats. Pharmacology Biochemistry and Behavior, 59, 105–108.

Modak, T., Bhad, R., & Rao, R. (2019) A rare case of physical dependence with psychedelic LSD - A case report. Journal of Substance Use, 24, 347–349.

Nagaeva, E., Zubarev, I., & Korpi, E. (2021) Electrophysiological Properties of Neurons: Current-Clamp Recordings in Mouse Brain Slices and Firing-Pattern Analysis. BIO-PROTOCOL, 11.

Nichols, D.E. (2016) Psychedelics. Pharmacological Reviews, 68, 264–355.

Nichols, D.E. & Frescas, S. (1999) Improvements to the Synthesis of Psilocybin and a Facile Method for Preparing the O-Acetyl Prodrug of Psilocin. Synthesis, 1999, 935–938.

Nutt, D., Spriggs, M., & Erritzoe, D. (2023) Psychedelics therapeutics: What we know, what we think, and what we need to research. Neuropharmacology, 223, 109257.

Parker, L.A. (1996) LSD produces place preference and flavor avoidance but does not produce flavor aversion in rats. Behavioral Neuroscience, 110, 503–508.

Pettit, H.O., Ettenberg, A., Bloom, F.E., & Koob, G.F. (1984) Destruction of dopamine in the nucleus accumbens selectively attenuates cocaine but not heroin self-administration in rats. Psychopharmacology, 84, 167–173.

Preller, K.H. & Vollenweider, F.X. (2018) Phenomenology, Structure, and Dynamic of Psychedelic States. In Halberstadt, A.L., Vollenweider, F.X., & Nichols, D.E. (eds), Behavioral Neurobiology of Psychedelic Drugs. Springer, Berlin, Heidelberg, pp. 221– 256.

Risinger, F.O., Dickinson, S.D., & Cunningham, C.L. (1992) Haloperidol reduces ethanol-induced motor activity stimulation but not conditioned place preference. Psychopharmacology, 107, 453–456.

Saal, D., Dong, Y., Bonci, A., & Malenka, R.C. (2003) Drugs of abuse and stress trigger a common synaptic adaptation in dopamine neurons. Neuron, 37, 577–582.

Sakloth, F., Leggett, E., Moerke, M.J., Townsend, E.A., Banks, M.L., & Negus, S.S. (2019) Effects of acute and repeated treatment with serotonin 5-HT2A receptor agonist hallucinogens on intracranial self-stimulation in rats. Experimental and Clinical Psychopharmacology, 27, 215–226.

Schlag, A.K., Aday, J., Salam, I., Neill, J.C., & Nutt, D.J. (2022) Adverse effects of psychedelics: From anecdotes and misinformation to systematic science. J Psychopharmacol, 36, 258–272.

Schmid, Y., Enzler, F., Gasser, P., Grouzmann, E., Preller, K.H., Vollenweider, F.X., Brenneisen, R., Müller, F., Borgwardt, S., & Liechti, M.E. (2015) Acute Effects of Lysergic Acid Diethylamide in Healthy Subjects. Biological Psychiatry, Serotonin, Mood, and Anxiety, 78, 544–553.

Shansky, R.M. & Murphy, A.Z. (2021) Considering sex as a biological variable will require a global shift in science culture. Nat Neurosci, 24, 457–464.

Siegel, R.K. & Jarvik, M.E. (1980) DMT self-administration by monkeys in isolation. Bull. Psychon. Soc., 16, 117–120.

Siivonen, M.S., de Miguel, E., Aaltio, J., Manner, A.K., Vahermo, M., Yli-Kauhaluoma, J., Linden, A.-M., Aitta-aho, T., & Korpi, E.R. (2018) Conditioned Reward of Opioids, but not Psychostimulants, is Impaired in GABA-A Receptor δ Subunit Knockout Mice. Basic & Clinical Pharmacology & Toxicology, 123, 558–566.

Stuber, G.D., Klanker, M., de Ridder, B., Bowers, M.S., Joosten, R.N., Feenstra, M.G., & Bonci, A. (2008) Reward-Predictive Cues Enhance Excitatory Synaptic Strength onto Midbrain Dopamine Neurons. Science, 321, 1690–1692.

Thomas, M.J. & Malenka, R.C. (2003) Synaptic plasticity in the mesolimbic dopamine system. Philosophical Transactions of the Royal Society of London. Series B: Biological Sciences, 358, 815–819.

Tzschentke, T.M. (1998) Measuring reward with the conditioned place preference paradigm: a comprehensive review of drug effects, recent progress and new issues. Progress in Neurobiology, 56, 613–672.

Vaccarino, F.J., Amalric, M., Swerdlow, N.R., & Koob, G.F. (1986) Blockade of amphetamine but not opiate-induced locomotion following antagonism of dopamine function in the rat. Pharmacology Biochemistry and Behavior, 24, 61–65.

Vargas-Perez, H., Minauro-Sanmiguel, F., Ting-A-Kee, R., Grieder, T.E., Méndez-Díaz, M., Prospéro-García, O., & van der Kooy, D. (2024) Rewarding Effects of the Hallucinogen 4-AcO-DMT Administration and Withdrawal in Rats: A Challenge to the Opponent-Process Theory. Neuroscience Letters, 820, 137597.

Vashchinkina, E., Manner, A.K., Vekovischeva, O., Hollander, B. den, Uusi-Oukari, M., Aitta-aho, T., & Korpi, E.R. (2014) Neurosteroid Agonist at GABAA Receptor Induces Persistent Neuroplasticity in VTA Dopamine Neurons. Neuropsychopharmacol, 39, 727–737.

Vashchinkina, E., Panhelainen, A., Vekovischeva, O.Yu., Aitta-aho, T., Ebert, B., Ator, N.A., & Korpi, E.R. (2012) GABA Site Agonist Gaboxadol Induces Addiction-Predicting Persistent Changes in Ventral Tegmental Area Dopamine Neurons But Is Not Rewarding in Mice or Baboons. J. Neurosci., 32, 5310–5320.

Vashchinkina, E., Piippo, O., Vekovischeva, O., Krupitsky, E., Ilyuk, R., Neznanov, N., Kazankov, K., Zaplatkin, I., & Korpi, E.R. (2018) Addiction-related interactions of pregabalin with morphine in mice and humans: reinforcing and inhibiting effects. Addiction Biology, 23, 945–958.

Volkow, N.D., Wise, R.A., & Baler, R. (2017) The dopamine motive system: implications for drug and food addiction. Nat Rev Neurosci, 18, 741–752.

Wise, R.A. & Bozarth, M.A. (1987) A psychomotor stimulant theory of addiction. Psychological Review, 94, 469–492.

Yanagita, T. (1986) Intravenous self-administration of (−)-cathinone and 2-amino-1-(2,5-dimethoxy-4-methyl)phenylpropane in rhesus monkeys. Drug and Alcohol Dependence, Committee on Problems of Drug Dependence Symposium on Stimulants and Hallucinogens, 17, 135–141.

