## Supplementary material for "Effects of serotonin agonists LSD and 25CN-NBOH on conditioned place preference and on synaptic plasticity of VTA dopamine neurons in mice"

for

### 1. Materials and reproducibility

**Table S1.** Key Resources Table

| Resource type | Designation | Source or reference | Identifier | Additional information |
|---|---|---|---|---|
| *Strain (Mus musculus)* | C57Bl/6JCrl | Charles River | MGI:3028467; RRID: IMSR_JAX:000664; CRL:632 | JAX® mice strain |
| *Genetic reagent (Mus musculus)* | Tg(Th-EGFP)DJ76Gsat/Mmnc | Gong et al. 2003 Nature PMID: 14586460 | RRID: MMRRC_000292-UNC | C57Bl/6J background |
| *Chemical compound, drug* | lysergic acid diethylamide (LSD) | Sigma-Aldrich/Merck | MERCK: L7007; CID: 5761 | |
| | lysergic acid diethylamide (LSD) | LGC Standards | LGC: FOR1346.0; CID:5761 | |
| | 25CN-NBOH hydrochloride | University of Copenhagen | CID: 90489020 | Gift from prof. Jesper L. Kristensen |
| | morphine hydrochloride | Yliopiston apteekki | CID: 5464110 | |
| *Software* | Activity Monitor | Med Associates | RRID: SCR_014296 | Version 6.02 |
| | EthoVision XT | Noldus | RRID: SCR_000441 | Version 10 |
| | Prism | GraphPad | RRID: SCR_002798 | Version 10.1 |
| | InVivoStat | Clark et al. 2012 J Psychopharmacol PMID: 22071578 | | Version 4.2 |
| *Apparatus* | Activity chambers | Med Associates | MA: ENV-51S | |
| | Place preference inserts | Med Associates | MA: ENV-517 | |
| | Multiclamp 700A microelectrode amplifier | Molecular Devices | RRID: SCR_018455 | |
| | Digidata 1440A Digitizer | Molecular Devices | RRID: SCR_021038 | |

#### 2. Methods: Bias assessment of the CPP setups

**Table S2.** Descriptive statistics of the distributions of times (s/min) spent on each of the floor materials used in the conditioned place preference experiments.

| **Unbiased** | N=124 | | | | 95%CI of mean | | |
|---|---|---|---|---|---|---|---|
| | Mean | Median | Min | Max | Lower | Upper | Std. dev. |
| Grid | 32.01 | 33.05 | 14.80 | 42.00 | 30.99 | 33.03 | 5.74 |
| Hole | 27.99 | 26.95 | 18.00 | 45.20 | 26.96 | 29.01 | 5.74 |
| **Biased** | N=64 | | | | 95%CI of mean | | |
| | Mean | Median | Min | Max | Lower | Upper | Std. dev. |
| Pink | 26.22 | 25.75 | 17.40 | 38.80 | 25.03 | 27.41 | 4.77 |
| Blue | 33.78 | 34.58 | 21.2 | 33.78 | 32.59 | 34.97 | 4.77 |
| *Males* | N=32 | | | | | | |
| Pink | 25.36 | 25.00 | 17.40 | 38.80 | 23.40 | 27.31 | 5.43 |
| Blue | 34.64 | 35.00 | 21.20 | 42.60 | 32.69 | 36.60 | 5.43 |
| *Females* | N=32 | | | | | | |
| Pink | 27.08 | 27.20 | 20.40 | 37.40 | 25.68 | 28.48 | 3.89 |
| Blue | 32.92 | 32.80 | 22.60 | 39.60 | 31.51 | 34.32 | 3.89 |

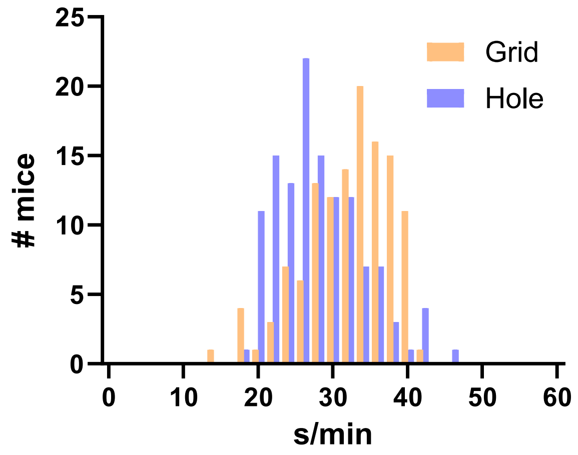

**Figure S1.** Frequency histograms of the distribution of the times spent on each floor material (mean s/min) during the pre-test sessions in the conditioned place preference experiments with the unbiased design (CPP1-4). N=124, all male mice.

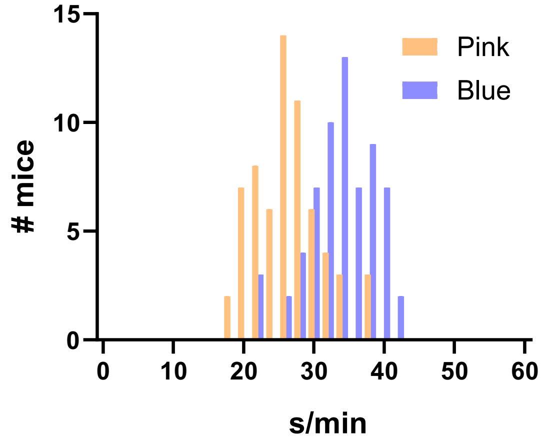

**Figure S2.** Frequency histograms of the distribution of the times spent on each floor material (mean s/min) during the pre-test sessions of the biased designs CPP5 and CPP6. N=64, combined data from all the mice.

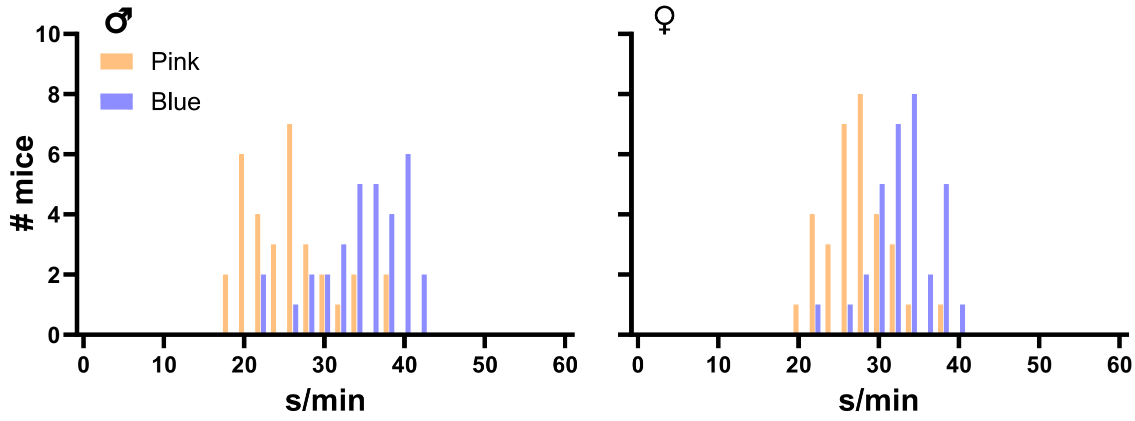

**Figure S3.** Frequency histograms of the distribution of the times spent on each floor material (mean s/min) during the pre-test sessions of the biased designs (CPP5 and CPP6) separated by the sex of the mice, male mice on the left, female mice on the right. N=32 on each.

#### 3. Methods: Error signal from the guillotine doors in CPP1–4

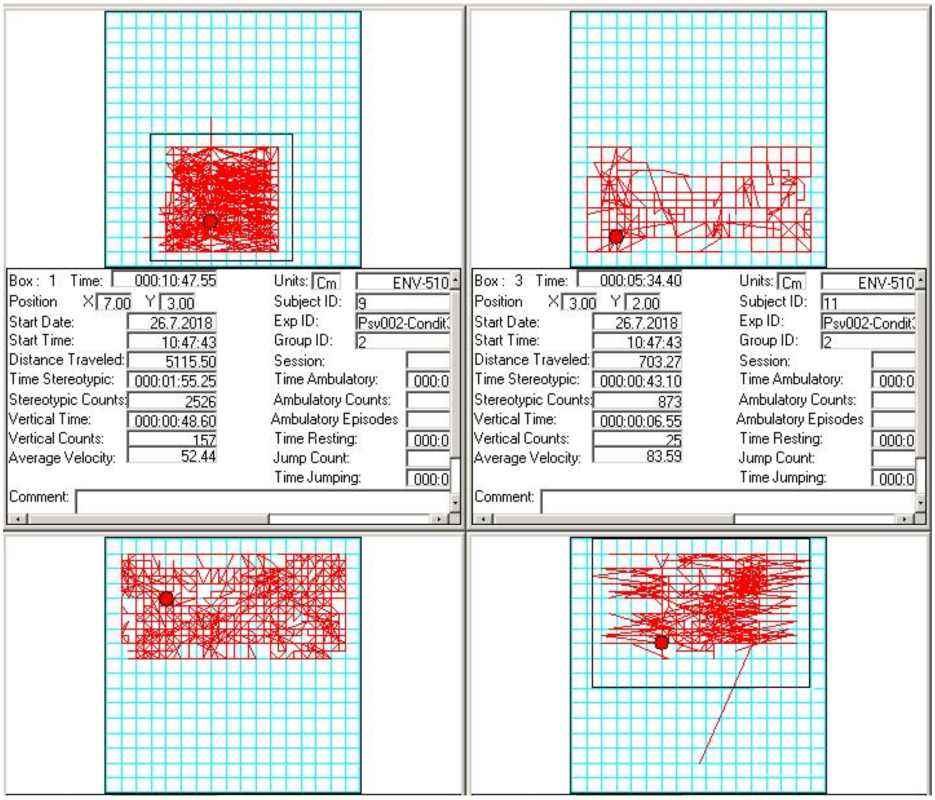

**Figure S4.** Closing of the guillotine door of the MedAssociates place preference inserts caused an error signal in some of the chambers by (presumably) interfering with the infrared beam at the site of the door, causing the recorded signal to constantly bounce between the actual location of the mouse and the door, causing the movement area of the mouse to be artificially restricted in the analysis software, as shown here in the screen capture of the Activity Monitor software (the upper left and lower right arenas).

#### 4. Results: The secondary analyses of the CPP experiments

**CPP1.** Similarly to the main analysis, the time-shift in the 25CN-NBOH group was larger (3.76 s/min [CI95% -1.1, 6.2]) than in the saline (1.8 s/min [CI95% -1.9, 5.6]) and the LSD-treated groups (1.07 [CI95% -2.5, 3.8]), but the permutation t-tests revealed no significant differences (all p>0.06; Figure 1DG).

**CPP2.** The secondary analysis of the time-shifts showed declines in all the groups (Saline -1.98 s/min [CI95% -4.1, 0.6], LSD -1.58 [CI95% -4.0, 0.55], 25CN-NBOH -1.72 s/min [CI95% -4.3, 0.26]) and none of the permutation t-test p-values reached the set level of significance (all p>0.1; Figure 1EH).

**CPP1+2**.The secondary analysis of the time-shifts showed neutral responses in all of the three groups (Saline 0.17 s/min [CI95% -1.9, 2.5], LSD 0.62 s/min [CI95% -2.7, 3.3], 25CN-NBOH 0.74 [CI95% -2.7, 3.3], Figure 1FI), reflected also in the results of the permutation t-tests (all p>0.5). These results disagree with the outcome of the main analysis by indicating no changes in preference in any of the treatment groups.

**CPP3**. The secondary analysis of the time-shift however showed a significant positive shift for the morphine-treated mice (4.41 s/min [CI95% 2.5, 6.5], permutation t-test p=0.02), but not for the saline (-2.11 s/min [CI95% -5.1, 1.0], permutation p=0.2) or the LSD groups (0.1 mg/kg 0.26 s/min [CI95% -3.0, 2.4], permutation p=0.8; 0.2 mg/kg 0.78 s/min [CI95% -2.4, 4.26], permutation p=0.8; Figure 2CE). Based on these results only morphine would have produced a significant preference.

**CPP4.** Also here, the secondary analysis of the time-shift (Figure 2DF) revealed a positive shift after the morphine treatment (3.3 s/min [CI95% 1.7, 6.2], permutation t-test p=0.01) and a slight decrease for the saline control group (-1.27 [CI95% -4.4, 1.0], permutation p=0.4). The lower dose of 25CN-NBOH tended to also cause a positive time-shift (1.5 mg/kg; 2.5 s/min [CI95% 0.7, 5.0], permutation p=0.07) whereas the shift was slightly negative for the higher dose (3.0 mg/kg; -1.65 s/min [CI95% -6.5, 1.4], permutation p=0.5). While not significant with the current numbers, the secondary analysis points towards a tendency for the lower dose of 25CN-NBOH to induce CPP.

**CPP5.** While in the main analysis none of the groups significantly differed fron one another, the secondary analysis contrasts this with the permutation t-tests for the within-group shifts showing all differences reaching the set level of statistical significance (all p<0.04; Figure 3E).

**CPP6.** Echoing the results of the main analysis, the secondary analysis with the permutation t-tests for the within-group shifts revealed differences reaching the level of significance in the morphine (p=0.02) and the lower 25CN-NBOH groups (1.5 mg/kg; p=0.03), but not in the saline (p=0.3) or in the higher 25CN-NBOH group (3.0 mg/kg; p=0.1).

#### CPP1 – unbiased design with LSD and 25CN-NBOH

##### Main analysis LSD + 25CN-NBOH vs saline control

| | Effect | Sums of squares | Degrees of freedom | Mean square | F-value | p | Sig. |
|---|---|---|---|---|---|---|---|
| Two-way ANOVA with Floor and Treatment as independent factors | floor | 123,49 | 1 | 123,49 | 11,30 | 0,0026 | ** |
| | treatment | 12,10 | 2 | 6,05 | 0,55 | 0,58 | |
| | floor * treatment | 80,30 | 2 | 40,15 | 3,68 | 0,041 | * |
| | Residual | 262,18 | 24 | 10,92 | | | |

| | Comparison | Difference | Lower 95% CI | Upper 95% CI | Std error | p | Sig. |
|---|---|---|---|---|---|---|---|
| All pairwise Floor*treatment comparisons with Holm's procedure | grid,lsd - hole,lsd | 3,587 | -0,728 | 7,901 | 2,09 | 0,92 | |
| | grid,lsd - grid,nboh | -1,427 | -5,741 | 2,888 | 2,09 | 1,00 | |
| | grid,lsd - hole,nboh | 6,853 | 2,539 | 11,168 | 2,09 | 0,04 | * |
| | grid,lsd - grid,saline | 3,187 | -1,128 | 7,501 | 2,09 | 0,92 | |
| | grid,lsd - hole,saline | 3,493 | -0,821 | 7,808 | 2,09 | 0,92 | |
| | hole,lsd - grid,nboh | -5,013 | -9,328 | -0,699 | 2,09 | 0,32 | |
| | hole,lsd - hole,nboh | 3,267 | -1,048 | 7,581 | 2,09 | 0,92 | |
| | hole,lsd - grid,saline | -0,4 | -4,714 | 3,914 | 2,09 | 1,00 | |
| | hole,lsd - hole,saline | -0,093 | -4,408 | 4,221 | 2,09 | 1,00 | |
| | grid,nboh - hole,nboh | 8,28 | 3,966 | 12,594 | 2,09 | 0,01 | ** |
| | grid,nboh - grid,saline | 4,613 | 0,299 | 8,928 | 2,09 | 0,41 | |
| | grid,nboh - hole,saline | 4,92 | 0,606 | 9,234 | 2,09 | 0,33 | |
| | hole,nboh - grid,saline | -3,667 | -7,981 | 0,648 | 2,09 | 0,92 | |
| | hole,nboh - hole,saline | -3,36 | -7,674 | 0,954 | 2,09 | 0,92 | |
| | grid,saline - hole,saline | 0,307 | -4,008 | 4,621 | 2,09 | 1,00 | |

##### Secondary analysis

| | Comparison | Difference | Lower 95% CI | Upper 95% CI | p | Sig. |
|---|---|---|---|---|---|---|
| Paired mean difference in Time on CS+ between the habituation and the test session and permutation t-test with 5000 resuhuffles | Saline | 1,80 | -1,88 | 5,48 | 0,40 | |
| | LSD 0.1 | 1,07 | -2,50 | 3,78 | 0,53 | |
| | 25CN-NBOH 3.0 | 3,76 | -1,06 | 6,17 | 0,06 | |

#### CPP2 – unbiased design with LSD and 25CN-NBOH

##### Main analysis LSD + 25CN-NBOH vs saline control

| | Effect | Sums of squares | Degrees of freedom | Mean square | F-value | p | Sig. |
|---|---|---|---|---|---|---|---|
| Two-way ANOVA with Floor and Treatment as independent factors | floor | 295,369 | 1 | 295,369 | 20,78 | 1E-04 | *** |
| | treatment | 1,308 | 2 | 0,654 | 0,05 | 0,955 | |
| | floor * treatment | 24,655 | 2 | 12,327 | 0,87 | 0,433 | |
| | Residual | 341,125 | 24 | 14,214 | | | |

| | Comparison | Difference | Lower 95% CI | Upper 95% CI | Std error | p | Sig. |
|---|---|---|---|---|---|---|---|
| All pairwise Floor*treatment comparisons with Holm's procedure | grid,lsd - hole,lsd | 7,907 | 2,985 | 12,828 | 2,384 | 0,043 | * |
| | grid,lsd - grid,nboh | 2,587 | -2,335 | 7,508 | 2,384 | 1,000 | |
| | grid,lsd - hole,nboh | 6,333 | 1,412 | 11,255 | 2,384 | 0,152 | |
| | grid,lsd - grid,saline | 0,56 | -4,361 | 5,481 | 2,384 | 1,000 | |
| | grid,lsd - hole,saline | 7,733 | 2,812 | 12,655 | 2,384 | 0,048 | * |
| | hole,lsd - grid,nboh | -5,32 | -10,241 | -0,399 | 2,384 | 0,318 | |
| | hole,lsd - hole,nboh | -1,573 | -6,495 | 3,348 | 2,384 | 1,000 | |
| | hole,lsd - grid,saline | -7,347 | -12,268 | -2,425 | 2,384 | 0,067 | |
| | hole,lsd - hole,saline | -0,173 | -5,095 | 4,748 | 2,384 | 1,000 | |
| | grid,nboh - hole,nboh | 3,747 | -1,175 | 8,668 | 2,384 | 0,904 | |
| | grid,nboh - grid,saline | -2,027 | -6,948 | 2,895 | 2,384 | 1,000 | |
| | grid,nboh - hole,saline | 5,147 | 0,225 | 10,068 | 2,384 | 0,329 | |
| | hole,nboh - grid,saline | -5,773 | -10,695 | -0,852 | 2,384 | 0,234 | |
| | hole,nboh - hole,saline | 1,4 | -3,521 | 6,321 | 2,384 | 1,000 | |
| | grid,saline - hole,saline | 7,173 | 2,252 | 12,095 | 2,384 | 0,073 | |

##### Secondary analysis

| | Comparison | Difference | Lower 95% CI | Upper 95% CI | p | Sig. |
|---|---|---|---|---|---|---|
| Paired mean difference in Time on CS+ between the habituation and the test session and permutation t-test with 5000 resuhuffles | Saline | -1,98 | -4,08 | 0,57 | 0,13 | |
| | LSD 0.1 | -1,58 | -4,03 | 0,55 | 0,25 | |
| | 25CN-NBOH 3.0 | -1,72 | -4,33 | 0,26 | 0,21 | |

#### CPP1+2 – unbiased designs with LSD and 25CN-NBOH combined

##### Main analysis LSD + 25CN-NBOH vs saline control

| | Effect | Sums of squares | Degrees of freedom | Mean square | F-value | p | Sig. |
|---|---|---|---|---|---|---|---|
| Two-way ANOVA with Floor and Treatment as independent factors and Experiment as a blocking factor | experiment | 899.646 | 1 | 899.646 | 66.59 | <0.0001 | *** |
| | floor | 400.417 | 1 | 400.417 | 29.64 | <0.0001 | *** |
| | treatment | 8.602 | 2 | 4.301 | 0.32 | 0.7287 | |
| | floor * treatment | 15.443 | 2 | 7.722 | 0.57 | 0.5681 | |
| | Residual | 716.071 | 53 | 13.511 | | | |

| | Comparison | Difference | Lower 95% CI | Upper 95% CI | Std error | p | Sig. |
|---|---|---|---|---|---|---|---|
| All pairwise Floor*treatment comparisons with Holm's procedure | grid,lsd - hole,lsd | 5.747 | 2.450 | 9.044 | 1.644 | 0.0125 | * |
| | grid,lsd - grid,nboh | 0.580 | -2.717 | 3.877 | 1.644 | 1.0000 | |
| | grid,lsd - hole,nboh | 6.593 | 3.296 | 9.890 | 1.644 | 0.0029 | ** |
| | grid,lsd - grid,saline | 1.873 | -1.424 | 5.170 | 1.644 | 1.0000 | |
| | grid,lsd - hole,saline | 5.613 | 2.316 | 8.910 | 1.644 | 0.0148 | * |
| | hole,lsd - grid,nboh | -5.167 | -8.464 | -1.870 | 1.644 | 0.0301 | * |
| | hole,lsd - hole,nboh | 0.847 | -2.450 | 4.144 | 1.644 | 1.0000 | |
| | hole,lsd - grid,saline | -3.873 | -7.170 | -0.576 | 1.644 | 0.1775 | |
| | hole,lsd - hole,saline | -0.133 | -3.430 | 3.164 | 1.644 | 1.0000 | |
| | grid,nboh - hole,nboh | 6.013 | 2.716 | 9.310 | 1.644 | 0.0082 | ** |
| | grid,nboh - grid,saline | 1.293 | -2.004 | 4.590 | 1.644 | 1.0000 | |
| | grid,nboh - hole,saline | 5.033 | 1.736 | 8.330 | 1.644 | 0.0345 | * |
| | hole,nboh - grid,saline | -4.720 | -8.017 | -1.423 | 1.644 | 0.0528 | |
| | hole,nboh - hole,saline | -0.980 | -4.277 | 2.317 | 1.644 | 1.0000 | |
| | grid,saline - hole,saline | 3.740 | 0.443 | 7.037 | 1.644 | 0.1888 | |

##### Secondary analysis

| | Comparison | Difference | Lower 95% CI | Upper 95% CI | p | Sig. |
|---|---|---|---|---|---|---|
| Paired mean difference in Time on CS+ between the habituation and the test session and permutation t-test with 5000 resuhuffles | Saline | 0,17 | -1,93 | 2,52 | 0,89 | |
| | LSD 0.1 | 0,62 | -1,27 | 2,41 | 0,52 | |
| | 25CN-NBOH 3.0 | 0,74 | -2,71 | 3,31 | 0,64 | |

#### CPP3 – unbiased design with LSD doses and morphine

##### Main analysis LSD vs saline control

| | Effect | Sums of squares | Degrees of freedom | Mean square | F-value | p | Sig. |
|---|---|---|---|---|---|---|---|
| Two-way ANOVA with Floor and Treatment as independent factors | floor | 5,7 | 1 | 5,7 | 0,2 | 0,7 | |
| | treatment | 2,0 | 2 | 1,0 | 0,0 | 1,0 | |
| | floor * treatment | 12,9 | 2 | 6,4 | 0,2 | 0,8 | |
| | Residual | 575,3 | 18 | 32,0 | | | |

| | Comparison | Difference | Lower 95% CI | Upper 95% CI | Std error | p | Sig. |
|---|---|---|---|---|---|---|---|
| All pairwise Floor*treatment comparisons with Holm's procedure | grid,lsd01 - hole,lsd01 | 2,83 | -5,6 | 11,2 | 4,0 | 1,0 | |
| | grid,lsd01 - grid,lsd02 | 1,58 | -6,8 | 10,0 | 4,0 | 1,0 | |
| | grid,lsd01 - hole,lsd02 | 2,43 | -6,0 | 10,8 | 4,0 | 1,0 | |
| | grid,lsd01 - grid,saline | 2,42 | -6,0 | 10,8 | 4,0 | 1,0 | |
| | grid,lsd01 - hole,saline | 1,67 | -6,7 | 10,1 | 4,0 | 1,0 | |
| | hole,lsd01 - grid,lsd02 | -1,25 | -9,6 | 7,1 | 4,0 | 1,0 | |
| | hole,lsd01 - hole,lsd02 | -0,40 | -8,8 | 8,0 | 4,0 | 1,0 | |
| | hole,lsd01 - grid,saline | -0,42 | -8,8 | 8,0 | 4,0 | 1,0 | |
| | hole,lsd01 - hole,saline | -1,17 | -9,6 | 7,2 | 4,0 | 1,0 | |
| | grid,lsd02 - hole,lsd02 | 0,85 | -7,5 | 9,2 | 4,0 | 1,0 | |
| | grid,lsd02 - grid,saline | 0,83 | -7,6 | 9,2 | 4,0 | 1,0 | |
| | grid,lsd02 - hole,saline | 0,08 | -8,3 | 8,5 | 4,0 | 1,0 | |
| | hole,lsd02 - grid,saline | -0,02 | -8,4 | 8,4 | 4,0 | 1,0 | |
| | hole,lsd02 - hole,saline | -0,77 | -9,2 | 7,6 | 4,0 | 1,0 | |
| | grid,saline - hole,saline | -0,75 | -9,1 | 7,6 | 4,0 | 1,0 | |

##### Positive control analysis morphine vs saline

| | Effect | Sums of squares | Degrees of freedom | Mean square | F-value | p | Sig. |
|---|---|---|---|---|---|---|---|
| Two-way ANOVA with Floor and Treatment as independent factors | floor | 9,3 | 1 | 9,3 | 0,2 | 0,6 | |
| | treatment | 12,8 | 1 | 12,8 | 0,3 | 0,6 | |
| | floor * treatment | 20,7 | 1 | 20,7 | 0,5 | 0,5 | |
| | Residual | 476,3 | 12 | 39,7 | | | |

| | Comparison | Difference | Lower 95% CI | Upper 95% CI | Std error | p | Sig. |
|---|---|---|---|---|---|---|---|
| All pairwise Floor*Treatment comparisons with Holm's procedure | grid,mor - hole,mor | 3,8 | -5,9 | 13,5 | 4,5 | 1,0 | |
| | grid,mor - grid,saline | 4,1 | -5,6 | 13,8 | 4,5 | 1,0 | |
| | grid,mor - hole,saline | 3,3 | -6,4 | 13,0 | 4,5 | 1,0 | |
| | hole,mor - grid,saline | 0,3 | -9,4 | 10,0 | 4,5 | 1,0 | |
| | hole,mor - hole,saline | -0,5 | -10,2 | 9,2 | 4,5 | 1,0 | |
| | grid,saline - hole,saline | -0,8 | -10,5 | 9,0 | 4,5 | 1,0 | |

#### Secondary analysis

| | Comparison | Difference | Lower 95% CI | Upper 95% CI | p | Sig. |
|---|---|---|---|---|---|---|
| Paired mean difference in Time on CS+ between the habituation and the test session and permutation t-test with 5000 resuhuffles | Saline | -2,1 | -5,1 | 1,0 | 0,23 | |
| | Morphine | 4,4 | 2,5 | 6,5 | 0,02 | * |
| | LSD 0.1 | 0,3 | -3,0 | 2,4 | 0,80 | |
| | LSD 0.2 | 0,8 | -2,4 | 4,3 | 0,69 | |

#### CPP4 – unbiased design with 25CN-NBOH doses and morphine

##### Main analysis 25CN-NBOH vs saline control

| | Effect | Sums of squares | Degrees of freedom | Mean square | F-value | p | Sig. |
|---|---|---|---|---|---|---|---|
| Two-way ANOVA with Floor and Treatment as independent factors | floor | 11,7 | 1 | 11,7 | 0,4 | 0,5 | |
| | treatment | 9,8 | 2 | 4,9 | 0,2 | 0,8 | |
| | floor * treatment | 80,2 | 2 | 40,1 | 1,4 | 0,3 | |
| | Residual | 519,5 | 18 | 28,9 | | | |

| | Comparison | Difference | Lower 95% CI | Upper 95% CI | Std error | p | Sig. |
|---|---|---|---|---|---|---|---|
| All pairwise Floor*treatment comparisons with Holm's procedure | grid,nboh15 - hole,nboh15 | 6,0 | -2,0 | 13,9 | 3,8 | 1,00 | |
| | grid,nboh15 - grid,nboh30 | 3,3 | -4,7 | 11,3 | 3,8 | 1,00 | |
| | grid,nboh15 - hole,nboh30 | 4,5 | -3,4 | 12,5 | 3,8 | 1,00 | |
| | grid,nboh15 - grid,saline | 3,9 | -4,1 | 11,8 | 3,8 | 1,00 | |
| | grid,nboh15 - hole,saline | 0,9 | -7,1 | 8,8 | 3,8 | 1,00 | |
| | hole,nboh15 - grid,nboh30 | -2,7 | -10,6 | 5,3 | 3,8 | 1,00 | |
| | hole,nboh15 - hole,nboh30 | -1,4 | -9,4 | 6,6 | 3,8 | 1,00 | |

| | Comparison | Difference | Lower 95% CI | Upper 95% CI | Std error | p | Sig. |
|---|---|---|---|---|---|---|---|
| | hole,nboh15 - grid,saline | -2,1 | -10,1 | 5,9 | 3,8 | 1,00 | |
| | hole,nboh15 - hole,saline | -5,1 | -13,1 | 2,9 | 3,8 | 1,00 | |
| | grid,nboh30 - hole,nboh30 | 1,2 | -6,7 | 9,2 | 3,8 | 1,00 | |
| | grid,nboh30 - grid,saline | 0,6 | -7,4 | 8,5 | 3,8 | 1,00 | |
| | grid,nboh30 - hole,saline | -2,4 | -10,4 | 5,5 | 3,8 | 1,00 | |
| | hole,nboh30 - grid,saline | -0,7 | -8,6 | 7,3 | 3,8 | 1,00 | |
| | hole,nboh30 - hole,saline | -3,7 | -11,6 | 4,3 | 3,8 | 1,00 | |
| | grid,saline - hole,saline | -3,0 | -11,0 | 5,0 | 3,8 | 1,00 | |

#### Positive control analysis morphine vs saline

| | Effect | Sums of squares | Degrees of freedom | Mean square | F-value | p | Sig. |
|---|---|---|---|---|---|---|---|
| Two-way ANOVA with Floor and Treatment as independent factors | floor | 48,8 | 1 | 48,8 | 2,7 | 0,1 | |
| | treatment | 4,3 | 1 | 4,3 | 0,2 | 0,6 | |
| | floor * treatment | 168,6 | 1 | 168,6 | 9,2 | 0,01 | * |
| | Residual | 219,1 | 12 | 18,3 | | | |

| | Comparison | Difference | Lower 95% CI | Upper 95% CI | Std error | p | Sig. |
|---|---|---|---|---|---|---|---|
| All pairwise Floor*treatment comparisons with Holm's procedure | grid,mor - hole,mor | 10,0 | 3,4 | 16,6 | 3,0 | 0,04 | * |
| | grid,mor - grid,saline | 5,5 | -1,1 | 12,0 | 3,0 | 0,39 | |
| | grid,mor - hole,saline | 2,5 | -4,1 | 9,0 | 3,0 | 0,68 | |
| | hole,mor - grid,saline | -4,5 | -11,1 | 2,1 | 3,0 | 0,48 | |
| | hole,mor - hole,saline | -7,5 | -14,1 | -1,0 | 3,0 | 0,14 | |
| | grid,saline - hole,saline | -3,0 | -9,6 | 3,6 | 3,0 | 0,68 | |

#### Secondary analysis

| | Comparison | Difference | Lower 95% CI | Upper 95% CI | p | Sig. |
|---|---|---|---|---|---|---|
| Paired mean difference in Time on CS+ between the habituation and the test session and permutation t-test with 5000 resuhuffles | Saline | -1,27 | -4,39 | 0,96 | 0,23 | |
| | Morphine | 3,30 | 1,72 | 6,23 | 0,01 | * |
| | 25CN-NBOH 1.5 | 2,51 | 0,71 | 4,98 | 0,07 | |
| | 25CN-NBOH 3.0 | 1,65 | -6,49 | 1,43 | 0,49 | |

#### CPP5 – biased design with LSD

##### Main analysis LSD vs saline control

| | Effect | Sums of squares | Degrees of freedom | Mean square | F-value | p | Sig. |
|---|---|---|---|---|---|---|---|
| One-way ANOVA with Treatment as an independent variable and Sex as a blocking factor | sex | 56,3 | 1 | 56,3 | 4,10 | 0,06 | |
| | treatment | 8,8 | 2 | 4,4 | 0,32 | 0,73 | |
| | residual | 274,6 | 20 | 13,7 | | | |

| | Comparison | Difference | Lower 95% CI | Upper 95% CI | Std error | p | Sig. |
|---|---|---|---|---|---|---|---|
| Pairwise comparisons against saline control with Holm's procedure | lsd01 - saline | -1,00 | -4,9 | 2,9 | 1,9 | 1,00 | |
| | lsd02 - saline | 0,45 | -3,4 | 4,3 | 1,9 | 1,00 | |

##### Positive control analysis morphine vs saline

| | Difference | Lower 95% CI | Upper 95% CI | t-statistic | Degrees c | p | Sig. |
|---|---|---|---|---|---|---|---|
| Unpaired t-test with Welch's correction | -2,3 | -6,2 | 1,5 | -1,3 | 11,8 | 0,21 | |

##### Secondary analysis

| | Comparison | Difference | Lower 95% CI | Upper 95% CI | p | Sig. |
|---|---|---|---|---|---|---|
| Time on CS+ between the pre-test and the test session and permutation t-test with 5000 resuhuffles | Saline | 4,73 | 2,82 | 6,36 | 0,0096 | ** |
| | Morphine | 7,08 | 4,62 | 10,1 | 1E-07 | *** |
| | LSD 0.1 | 3,76 | 1 | 6,99 | 0,0414 | * |
| | LSD 0.2 | 5,19 | 1,66 | 7,45 | 0,0226 | * |

#### CPP6 – biased design with 25CN-NBOH

##### Main analysis LSD vs saline control

| | Effect | Sums of squares | Degrees of freedom | Mean square | F-value | p | Sig. |
|---|---|---|---|---|---|---|---|
| One-way ANOVA with Treatment as an independent variable and Sex as a blocking factor | sex | 14,0 | 1 | 14,0 | 0,53 | 0,48 | |
| | treatment | 237,6 | 2 | 118,8 | 4,48 | 0,02 | * |
| | residual | 530,0 | 20 | 26,5 | | | |

| | Comparison | Difference | Lower 95% CI | Upper 95% CI | Std error | p | Sig. |
|---|---|---|---|---|---|---|---|
| Pairwise comparisons against saline control with Holm's procedure | nboh1.5 - saline | 7,68 | 2,3 | 13,1 | 2,6 | 0,01 | * |
| | nboh3.0 - saline | 4,356 | -1,01 | 9,7 | 2,6 | 0,11 | |

##### Positive control analysis morphine vs saline

| | Difference | Lower 95% CI | Upper 95% CI | t-statistic | Degrees of freedom | p | Sig. |
|---|---|---|---|---|---|---|---|
| Unpaired t-test with Welch's correction | -7,1 | -12,5 | -2,0 | -2,8 | 14,0 | 0,02 | * |

##### Secondary analysis

| | Comparison | Difference | Lower 95% CI | Upper 95% CI | p | Sig. |
|---|---|---|---|---|---|---|
| Time on CS+ between the habituation and the test session and permutation t-test with 5000 resuhuffles | Saline | -1,75 | -4,81 | 1,60 | 0,337 | |
| | Morphine | 5,3 | 2,16 | 8,75 | 0,019 | * |
| | 25CN-NBOH 1.5 | 5,98 | 2,56 | 10 | 0,0258 | * |
| | 25CN-NBOH 3.0 | 2,63 | -0,188 | 5,5 | 0,132 | |

#### CPP5 - biased design with LSD

##### Positive control analysis against morphine

| CS– trials | | df | F | p | Sig. |
|---|---|---|---|---|---|
| Two-way repeated measures mixed model analysis | sex | 1,13 | 1,1 | 0,32 | |
| | treatment | 1,13 | 0,0 | 0,97 | |
| | session | 3,42 | 0,9 | 0,44 | |
| | treatmen*session | 3,42 | 2,7 | 0,06 | |

| Multiple comparisons within each session | | | Mean diff | 95% CI lower | 95% CI upper | p (un-adjusted) | p (Holm) | Sig. |
|---|---|---|---|---|---|---|---|---|
| 1 | Mor – | Sal | -9,3 | -21,3 | 2,7 | 0,12 | 0,48 | |
| 2 | Mor – | Sal | -1,5 | -13,5 | 10,5 | 0,80 | 0,98 | |
| 3 | Mor – | Sal | 4,1 | -7,9 | 16,1 | 0,49 | 0,98 | |
| 4 | Mor – | Sal | 7,4 | -4,6 | 19,4 | 0,22 | 0,66 | |

| CS+ trials | | df | F | p | Sig. |
|---|---|---|---|---|---|
| Two-way repeated measures mixed model analysis | sex | 1,13 | 0,4 | 0,56 | |
| | treatment | 1,13 | 192,2 | <0.0001 | *** |
| | session | 3,42 | 2,3 | 0,08 | |
| | treatmen*session | 3,42 | 0,7 | 0,54 | |

| Multiple comparisons within each session | | | Mean diff | 95% CI lower | 95% CI upper | p (un-adjusted) | p (Holm) | Sig. |
|---|---|---|---|---|---|---|---|---|
| 1 | Mor – | Sal | 143,2 | 108,1 | 178,4 | 0,00 | 0,00 | *** |
| 2 | Mor – | Sal | 173,7 | 138,6 | 208,9 | 0,00 | 0,00 | *** |
| 3 | Mor – | Sal | 151,6 | 116,4 | 186,7 | 0,00 | 0,00 | *** |
| 4 | Mor – | Sal | 160,8 | 125,7 | 195,9 | 0,00 | 0,00 | *** |

##### Primary analysis of LSD against saline

| CS– trials | | df | F | p | Sig. |
|---|---|---|---|---|---|
| Two-way repeated measures mixed model analysis | sex | 1, 20 | 1,4 | 0,24 | |
| | treatment | 2, 20 | 0,6 | 0,55 | |
| | session | 3, 63 | 6,4 | 0,00 | *** |
| | treatment*session | 6, 63 | 0,8 | 0,60 | |

| Multiple comparisons within each session | | | Mean diff | 95% CI lower | 95% CI upper | p (un-adjusted) | p (Holm) | Sig. |
|---|---|---|---|---|---|---|---|---|
| 1 | LSD0.1 – | Sal | 7,9 | -4,2 | 19,96 | 0,20 | 1,00 | |
| | | LSD0.2 | 2,3 | -9,76 | 14,38 | 0,70 | 1,00 | |
| | LSD0.2 – | Sal | 5,6 | -6,49 | 17,65 | 0,36 | 1,00 | |
| 2 | LSD0.1 – | Sal | -4,5 | -16,57 | 7,57 | 0,46 | 1,00 | |
| | | LSD0.2 | -5,6 | -17,65 | 6,5 | 0,36 | 1,00 | |
| | LSD0.2 – | Sal | 1,1 | -10,9 | 13,2 | 0,86 | 1,00 | |

| | | | Mean diff | 95% CI lower | 95% CI upper | p (un-adjusted) | p (Holm) | Sig. |
|---|---|---|---|---|---|---|---|---|
| 3 | LSD0.1 – | Sal | 5,3 | -6,8 | 17,3 | 0,39 | 1,00 | |
| | | LSD0.2 | 1,3 | -10,8 | 13,4 | 0,86 | 1,00 | |
| | LSD0.2 – | Sal | 4,0 | -8,1 | 16 | 0,51 | 1,00 | |
| 4 | LSD0.1 – | Sal | 6,1 | -6 | 18,2 | 0,32 | 1,00 | |
| | | LSD0.2 | -1,9 | -14 | 10,2 | 0,75 | 1,00 | |
| | LSD0.2 – | Sal | 8,0 | -4,1 | 20,1 | 0,19 | 1,00 | |

| CS+ trials | | df | F | p | Sig. |
|---|---|---|---|---|---|
| Two-way repeated measures mixed model analysis | sex | 1, 20 | 0,4 | 0,56 | |
| | treatment | 2, 20 | 1,2 | 0,33 | |
| | session | 3, 63 | 3,2 | 0,03 | * |
| | treatment*session | 6, 63 | 0,5 | 0,80 | |

| Multiple comparisons within each session | | | Mean diff | 95% CI lower | 95% CI upper | p (un-adjusted) | p (Holm) | Sig. |
|---|---|---|---|---|---|---|---|---|
| 1 | LSD0.1 – | Sal | 2,1 | -14,6 | 18,9 | 0,32 | 1,00 | |
| | | LSD0.2 | 8,3 | -8,4 | 25,1 | 0,32 | 1,00 | |
| | LSD0.2 – | Sal | -6,2 | -23 | 10,6 | 0,46 | 1,00 | |
| 2 | LSD0.1 – | Sal | -6,5 | -23,3 | 10,2 | 0,43 | 1,00 | |
| | | LSD0.2 | 8,9 | -7,9 | 25,6 | 0,30 | 1,00 | |
| | LSD0.2 – | Sal | -15,4 | -32,2 | 1,4 | 0,07 | 0,84 | |
| 3 | LSD0.1 – | Sal | -2,3 | -19,1 | 14,5 | 0,78 | 1,00 | |
| | | LSD0.2 | 0,6 | -16,1 | 17,4 | 0,94 | 1,00 | |
| | LSD0.2 – | Sal | -2,9 | -19,7 | 13,8 | 0,73 | 1,00 | |
| 4 | LSD0.1 – | Sal | -8,6 | -25,4 | 8,2 | 0,31 | 1,00 | |
| | | LSD0.2 | 3,4 | -13,4 | 20,2 | 0,69 | 1,00 | |
| | LSD0.2 – | Sal | -11,9 | -28,7 | 4,8 | 0,16 | 1,00 | |

#### CPP6 - Biased design with 25CN-NBOH

##### Positive control analysis against morphine

| CS– trials | | df | F | p | Sig. |
|---|---|---|---|---|---|
| Two-way repeated measures mixed model analysis | sex | 1,13 | 0,1 | 0,75 | |
| | treatment | 1,13 | 1,6 | 0,23 | |
| | session | 2,42 | 4,3 | 0,01 | ** |
| | treatment*session | 3,42 | 0,9 | 0,47 | |

| Multiple comparisons within each session | | | Mean diff | 95% CI lower | 95% CI upper | p (un-adjusted) | p (Holm) | Sig. |
|---|---|---|---|---|---|---|---|---|
| 1 | Mor – | Sal | -4,6 | -20,3 | 11,0 | 0,55 | 1 | |
| 2 | Mor – | Sal | -12,4 | -28,1 | 3,2 | 0,12 | 0,48 | |
| 3 | Mor – | Sal | -11,5 | -27,2 | 4,1 | 0,14 | 0,48 | |
| 4 | Mor – | Sal | -1,8 | -17,4 | 13,9 | 0,82 | 1 | |

| CS+ trials | | df | F | p | Sig. |
|---|---|---|---|---|---|
| Two-way repeated measures mixed model analysis | sex | 1,13 | 0,1 | 0,83 | |
| | treatment | 1,13 | 32,5 | <0.0001 | *** |
| | session | 3,42 | 3,0 | 0,04 | * |
| | treatmen*session | 3,42 | 2,5 | 0,08 | |

| Multiple comparisons within each session | | | Mean diff | 95% CI lower | 95% CI upper | p (un-adjusted) | p (Holm) | Sig. |
|---|---|---|---|---|---|---|---|---|
| 1 | Mor – | Sal | 65,3 | 24,1 | 106,4 | 0,00 | 0,03 | ** |
| 2 | Mor – | Sal | 110,9 | 69,7 | 152,0 | < 0.0001 | 0,00 | *** |
| 3 | Mor – | Sal | 106,3 | 65,2 | 147,4 | < 0.0001 | 0,00 | *** |
| 4 | Mor – | Sal | 101,2 | 60,1 | 142,3 | < 0.0001 | 0,00 | *** |

**Primary analysis of 25CN-NBOH against saline**

| CS– trials | | df | F | p | Sig. |
|---|---|---|---|---|---|
| Two-way repeated measures mixed model analysis | sex | 1, 20 | 0,0 | 0,84 | |
| | treatment | 2, 20 | 2,5 | 0,11 | |
| | session | 3, 63 | 1,5 | 0,23 | |
| | treatment*session | 6, 63 | 0,4 | 0,90 | |

| Multiple comparisons within each session | | | Mean diff | 95% CI lower | 95% CI upper | p (un-adjusted) | p (Holm) | Sig. |
|---|---|---|---|---|---|---|---|---|
| 1 | 25CN-NBOH1.5 – | Sal | -10,3 | -26,2 | 5,6 | 0,19 | 1,00 | |
| | | 25CN-NBOH3.0 | -11,3 | -27,2 | 4,6 | 0,16 | 1,00 | |
| | 25CN-NBOH3.0 – | Sal | 1,0 | -149 | 16,9 | 0,90 | 1,00 | |
| 2 | 25CN-NBOH1.5 – | Sal | -12,6 | -28,5 | 3,3 | 0,12 | 1,00 | |
| | | 25CN-NBOH3.0 | -14,3 | -30,2 | 1,6 | 0,08 | 0,96 | |
| | 25CN-NBOH3.0 – | Sal | 1,7 | -14,2 | 17,6 | 0,83 | 1,00 | |
| 3 | 25CN-NBOH1.5 – | Sal | -3,9 | -19,8 | 12 | 0,62 | 1,00 | |
| | | 25CN-NBOH3.0 | -13,3 | -29,2 | 2,6 | 0,10 | 1,00 | |
| | 25CN-NBOH3.0 – | Sal | 9,4 | -6,5 | 25,3 | 0,24 | 1,00 | |
| 4 | 25CN-NBOH1.5 – | Sal | -4,2 | -20,1 | 11,7 | 0,60 | 1,00 | |
| | | 25CN-NBOH3.0 | -13,1 | -29 | 2,8 | 0,60 | 1,00 | |
| | 25CN-NBOH3.0 – | Sal | 8,9 | -7 | 24,8 | 0,27 | 1,00 | |

| CS+ trials | | df | F | p | Sig. |
|---|---|---|---|---|---|
| Two-way repeated measures mixed model analysis | sex | 1, 20 | 0,4 | 0,54 | |
| | treatment | 2, 20 | 3,6 | 0,05 | * |
| | session | 3, 63 | 3,3 | 0,03 | * |
| | treatment*session | 6, 63 | 1,1 | 0,37 | |

| Multiple comparisons within each session | | | Mean diff | 95% CI lower | 95% CI upper | p (un-adjusted) | p (Holm) | Sig. |
|---|---|---|---|---|---|---|---|---|
| 1 | 25CN-NBOH1.5 – | Sal | -7,9 | -27,1 | 11,4 | 0,42 | 1,00 | |
| | | 25CN-NBOH3.0 | 6,5 | -12,7 | 25,7 | 0,50 | 1,00 | |
| | 25CN-NBOH3.0 – | Sal | -24,4 | -33,6 | 4,8 | 0,14 | 1,00 | |
| 2 | 25CN-NBOH1.5 – | Sal | -16,9 | -36,2 | 2,2 | 0,08 | 0,72 | |
| | | 25CN-NBOH3.0 | -2,6 | -21,8 | 16,6 | 0,80 | 1,00 | |
| | 25CN-NBOH3.0 – | Sal | -5,6 | -24,8 | 13,6 | 0,56 | 0,84 | |
| 3 | 25CN-NBOH1.5 – | Sal | -18,9 | -38,2 | 0,3 | 0,05 | 0,50 | |
| | | 25CN-NBOH3.0 | -5,6 | -24,8 | 13,6 | 0,56 | 1,00 | |
| | 25CN-NBOH3.0 – | Sal | -13,3 | -32,6 | 5,9 | 0,17 | 1,00 | |
| 4 | 25CN-NBOH1.5 – | Sal | -27,3 | -46,5 | -8,1 | 0,01 | 0,07 | |
| | | 25CN-NBOH3.0 | -20,7 | -39,9 | -1,5 | 0,04 | 0,40 | |
| | 25CN-NBOH3.0 – | Sal | -6,6 | -25,8 | 12,6 | 0,49 | 1,00 | |

#### VTA dopamine AMPA:NMDA - average

**Main analysis LSD + 25CN-NBOH vs saline**

| | Effect | Sums of squares | Degrees of freedom | Mean square | F-value | p | Sig. |
|---|---|---|---|---|---|---|---|
| One-way ANOVA with Treatment as Independent variable and Sex as a blocking factor | sex | 0,001 | 1 | 0,001 | 0,01 | 0,91 | |
| | treatment | 0,8 | 2 | 0,4 | 6,09 | 0,0062 | ** |
| | Residual | 1,9 | 29 | 0,07 | | | |

| | Comparison | Difference | Lower 95% CI | Upper 95% CI | Std error | p | Sig. |
|---|---|---|---|---|---|---|---|
| Pairwise comparisons against saline control with Holm's procedure | lsd - saline | 0,36 | 0,15 | 0,57 | 0,10 | 0,0032 | ** |
| | nboh - saline | 0,16 | -0,082 | 0,41 | 0,12 | 0,18 | |

#### Positive control analysis morphine vs saline

| | Difference | Lower 95% CI | Upper 95% CI | t-statistic | Degrees of freedom | p | Sig. |
|---|---|---|---|---|---|---|---|
| Unpaired t-test | -0,3 | -0,5 | 0,0 | -2,5 | 17,0 | 0,02 | * |

**Secondary analysis**

| | Comparison | Difference | Lower 95% CI | Upper 95% CI | p | Sig. |
|---|---|---|---|---|---|---|
| Unpaired mean difference in AMPA:NMDA ratio between the drug and Saline control and a permutation t-test with 5000 reshuffles | Morphine | 0,289 | 0,00843 | 0,585 | 0,0222 | * |
| | LSD 0.1 | 0,414 | 0,152 | 0,853 | 0,0026 | ** |
| | 25CN-NBOH 3 | 0,114 | -0,0225 | 0,327 | 0,19 | |

#### VTA dopamine AMPA:NMDA - lateral cell population

**Main analysis LSD + 25CN-NBOH vs saline**

| | Effect | Sums of squares | Degrees of freedom | Mean square | F-value | p | Sig. |
|---|---|---|---|---|---|---|---|
| One-way ANOVA with Treatment as Independent variable and Sex as a blocking factor | sex | 0 | 1 | 0 | 0 | 1,00 | |
| | treatment | 0,02 | 2 | 0,01 | 0,14 | 0,87 | |
| | Residual | 1,64 | 22 | 0,07 | | | |

| | Comparison | Difference | Lower 95% CI | Upper 95% CI | Std error | p | Sig. |
|---|---|---|---|---|---|---|---|
| Pairwise comparisons against saline control with Holm's procedure | lsd - saline | 0,07 | -0,21 | 0,35 | 0,14 | 1,0 | |
| | nboh - saline | 0,001 | -0,27 | 0,27 | 0,13 | 1,0 | |

**Secondary analysis**

| | Comparison | Difference | Lower 95% CI | Upper 95% CI | p | Sig. |
|---|---|---|---|---|---|---|
| Unpaired mean difference in AMPA:NMDA ratio between the drug and Saline control and a permutation t-test with 5000 reshuffles | LSD 0.1 | 0,025 | -0,138 | 0,184 | 0,805 | |
| | 25CN-NBOH 3.0 | -0,0321 | -0,184 | 0,0716 | 0,724 | |

#### VTA dopamine AMPA:NMDA - medial cell population

**Main analysis LSD + 25CN-NBOH vs saline**

| | Effect | Sums of squares | Degrees of freedom | Mean square | F-value | p | Sig. |
|---|---|---|---|---|---|---|---|
| One-way ANOVA with Treatment as Independent variable and Sex as a blocking factor | sex | 0,01 | 1 | 0,01 | 0,09 | 0,77 | |
| | treatment | 1,10 | 2 | 0,55 | 6,49 | 0,0050 | ** |
| | residual | 2,28 | 27 | 0,08 | | | |

| | Comparison | Difference | Lower 95% CI | Upper 95% CI | Std error | p | Sig. |
|---|---|---|---|---|---|---|---|
| Pairwise comparisons against saline control with Holm's procedure | lsd - saline | 0,415 | 0,175 | 0,656 | 0,117 | 0,0029 | ** |
| | nboh - saline | 0,267 | -0,026 | 0,56 | 0,143 | 0,0728 | |

**Secondary analysis**

| | Comparison | Difference | Lower 95% CI | Upper 95% CI | p | Sig. |
|---|---|---|---|---|---|---|
| Unpaired mean difference in AMPA:NMDA ratio between the drug and Saline control and a permutation t-test with 5000 reshuffles | LSD 0.1 | 0,468 | 0,184 | 0,874 | 0,0036 | ** |
| | 25CN-NBOH 3 | 0,297 | 0,00933 | 0,745 | 0,0512 | |

**VTA dopamine AMPA:NMDA - average (1–2 cells/animal)**

| | mean | 95%CI lower | 95%CI upper | median | s | n |
|---|---|---|---|---|---|---|
| saline | 0,283 | 0,192 | 0,374 | 0,249 | 0,165 | 15 |
| LSD 0.1 | 0,697 | 0,32 | 1,075 | 0,514 | 0,562 | 11 |
| 25CN-NBOH 3.0 | 0,398 | 0,195 | 0,6 | 0,337 | 0,219 | 7 |
| Morphine 10 | 0,572 | 0,046 | 1,097 | 0,55 | 0,33 | 4 |

Unpaired mean difference and two-sided permutation t-test *vs* saline (bootstrapped with 5000 resufles of control and test labels)

| ***vs. saline*** | mean diff | 95%CI lower | 95%CI upper | p |
|---|---|---|---|---|
| LSD 0.1 | 0,414 | 0,152 | 0,853 | 0,0026 |
| 25CN-NBOH 3.0 | 0,114 | -0,0225 | 0,327 | 0,19 |
| Morphine 10 | 0,289 | 0,00843 | 0,585 | 0,0222 |

**VTA dopamine AMPA:NMDA - lateral cell population**

| | mean | 95%CI lower | 95%CI upper | median | s | n |
|---|---|---|---|---|---|---|
| saline | 0,292 | 0,177 | 0,406 | 0,28 | 0,205 | 13 |
| LSD 0.1 | 0,313 | 0,145 | 0,481 | 0,295 | 0,17 | 6 |
| 25CN-NBOH 3.0 | 0,257 | 0,116 | 0,398 | 0,24 | 0,102 | 7 |
| Morphine 10 | 0,52 | 0,2 | 0,84 | 0,52 | 0,255 | 2 |

unpaired mean difference and two-sided permutation t-test vs saline (bootstrapped with 5000 resufles of control and test labels)

| ***vs. saline*** | mean diff | 95%CI lower | 95%CI upper | p |
|---|---|---|---|---|
| LSD 0.1 | 0,025 | -0,138 | 0,184 | 0,805 |
| 25CN-NBOH 3.0 | -0,0321 | -0,184 | 0,0716 | 0,724 |

**VTA dopamine AMPA:NMDA - medial cell population**

| | mean | 95%CI lower | 95%CI upper | median | s | n |
|---|---|---|---|---|---|---|
| saline | 0,292 | 0,0678 | 0,516 | 0,23 | 0,227 | 14 |
| LSD 0.1 | 0,761 | 0,5078 | 1,014 | 0,52 | 0,558 | 11 |
| 25CN-NBOH 3.0 | 0,59 | 0,311 | 0,869 | 0,35 | 0,496 | 6 |
| Morphine 10 | 0,897 | 0,569 | 1,225 | 1,07 | 0,446 | 3 |

unpaired mean difference and two-sided permutation t-test vs saline (bootstrapped with 5000 resufles of control and test labels)

| ***vs. saline*** | mean diff | 95%CI lower | 95%CI upper | p |
|---|---|---|---|---|
| LSD 0.1 | 0,468 | 0,184 | 0,874 | 0,0036 |
| 25CN-NBOH 3.0 | 0,297 | -0,00933 | 0,745 | 0,0512 |
